# An implicit memory of errors limits human sensorimotor adaptation

**DOI:** 10.1101/868406

**Authors:** Scott T. Albert, Jihoon Jang, Hannah Sheahan, Lonneke Teunissen, Koenraad Vandevoorde, Reza Shadmehr

## Abstract

After extended practice, motor adaptation reaches a limit in which learning appears to stop, despite the fact that residual errors persist. What prevents the brain from eliminating the residual errors? Here we found that the adaptation limit was causally dependent on the second order statistics of the perturbation; when variance was high, learning was impaired and large residual errors persisted. However, when learning relied solely on explicit strategy, both the adaptation limit and its dependence on perturbation variability disappeared. In contrast, when learning depended entirely, or in part on implicit learning, residual errors developed. Residual errors in implicit performance were caused by variance-dependent modifications to error sensitivity, not forgetting. These observations are consisted with a model of learning in which the implicit system becomes more sensitive to error when errors are consistent, but forgets this memory of errors over time. Thus, residual errors in motor adaptation are a signature of the implicit learning system, caused by an error sensitivity that depends on the history of past errors.

## Introduction

During motor adaptation, perturbations alter the sensory consequences of motor commands, yielding sensory prediction errors. In humans and other animals, the brain learns from these errors and adjusts its motor commands on subsequent attempts. Over many trials, the adjustments accumulate, but surprisingly, adaptation often remains incomplete; even after extended periods of practice, residual errors persist in many behaviors including reaching^1–4^, saccades^5,6^, and walking^7^. Why does learning appear to stop despite the fact that errors remain?

Current models suggest that adaptation is supported by distinct learning systems: one implicit^8^, and the other explicit^9–11^. It is thought that the implicit system contributes little to modulation of asymptotic performance; when challenged with fixed errors, the implicit system appears to saturate at identical levels^12,13^. In contrast, explicit strategy provides greater flexibility; its asymptotic behavior is altered as people age^14–16^, under different types of feedback^17^, and with the time allotted for the preparation of a movement^18^. Therefore, current evidence suggests that the explicit system alone modifies the asymptotic state of learning.

Here we tested this view using stochastic perturbations that affected reaching movements. We found that when perturbation variability was high, residual errors increased^19–21^. Furthermore, when perturbation variability was increased mid-experiment, the asymptotic performance decreased, causing participants to lose what they had already learned. Thus, the asymptote of adaptation was not a hard limit, but a dynamic variable that depended on the second order statistics of the perturbation. Which adaptive system was responsible for limiting the adaptation process?

To answer this question we isolated implicit and explicit components of adaptation using several methodologies including verbal instructions, aim reporting^9^, limiting reaction time^22–24^, and delaying visual feedback^25,26^. Across all experiments there was a very consistent pattern; learning that relied solely on explicit strategy did not suffer from residual errors, and was not affected by perturbation variance. That is, in contrast to earlier findings, the asymptotic limit of adaptation was due to an inherent property of the implicit system, a property that depended on perturbation variance.

Why did the implicit system suffer from an inability to eliminate residual errors, and why did the impairment become greater when perturbation variance increased? Implicit adaptation is supported by two competing processes, learning and forgetting^27–30^. Learning is controlled by sensitivity to error, and forgetting is controlled by the rate at which memory decays over time. When errors are large, learning dominates, yielding changes in motor commands that improve performance. However, as errors get small, forgetting reaches an equilibrium with error-based learning. Thus, in theory the asymptote of implicit adaptation could be a result of an equilibrium between forces that promote learning, and forces that promote forgetting^17^. Changes to either of these underlying processes could in principle alter the total extent of implicit adaptation.

By measuring patterns of forgetting and trial-by-trial learning, we found that changes in the asymptote of implicit learning were achieved solely through modulation of its error sensitivity. Furthermore, the spatiotemporal properties of error sensitivity suggested that the brain updated its implicit learning processes according to the sequence of past errors^31^. When errors of a particular size were consistent, the brain increased its sensitivity to those errors. Furthermore, like adapted behavior, this memory of error consistency appeared to be limited by decay. The resulting model of implicit learning accounted not only for changes in the asymptotic extent of adaptation induced by stochastic perturbations^19–21^, but also the saturation of learning under error-clamp conditions^6^, and the dissolution of savings over time^32,33^. Overall, we report that the asymptotic limit of motor adaptation has a simple cause; the implicit system has an error sensitivity that is modulated by the history of past errors.

## Results

In an earlier study, Fernandes and colleagues^19^ exposed participants to variable visuomotor rotations (Fig. 1A, Rotation). All groups were exposed to a sequence of perturbations that had the same mean (30°), but different amounts of variability; one group experienced a constant perturbation of 30° (zero variance), while the other two groups experienced perturbations with low variance or high variance (Fig. 1B, top). At the end of training, reach angles in each group had saturated, but still yielded persistent residual errors. Curiously, residual errors increased with the variance of the perturbation (Fig. 1H, Fernandes, median residual error on last 10 trials; repeated measures ANOVA: F(2,14)=17.8, p<0.001, 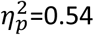). Why did perturbation variance reduce the total extent of adaptation?

**Figure 1.**
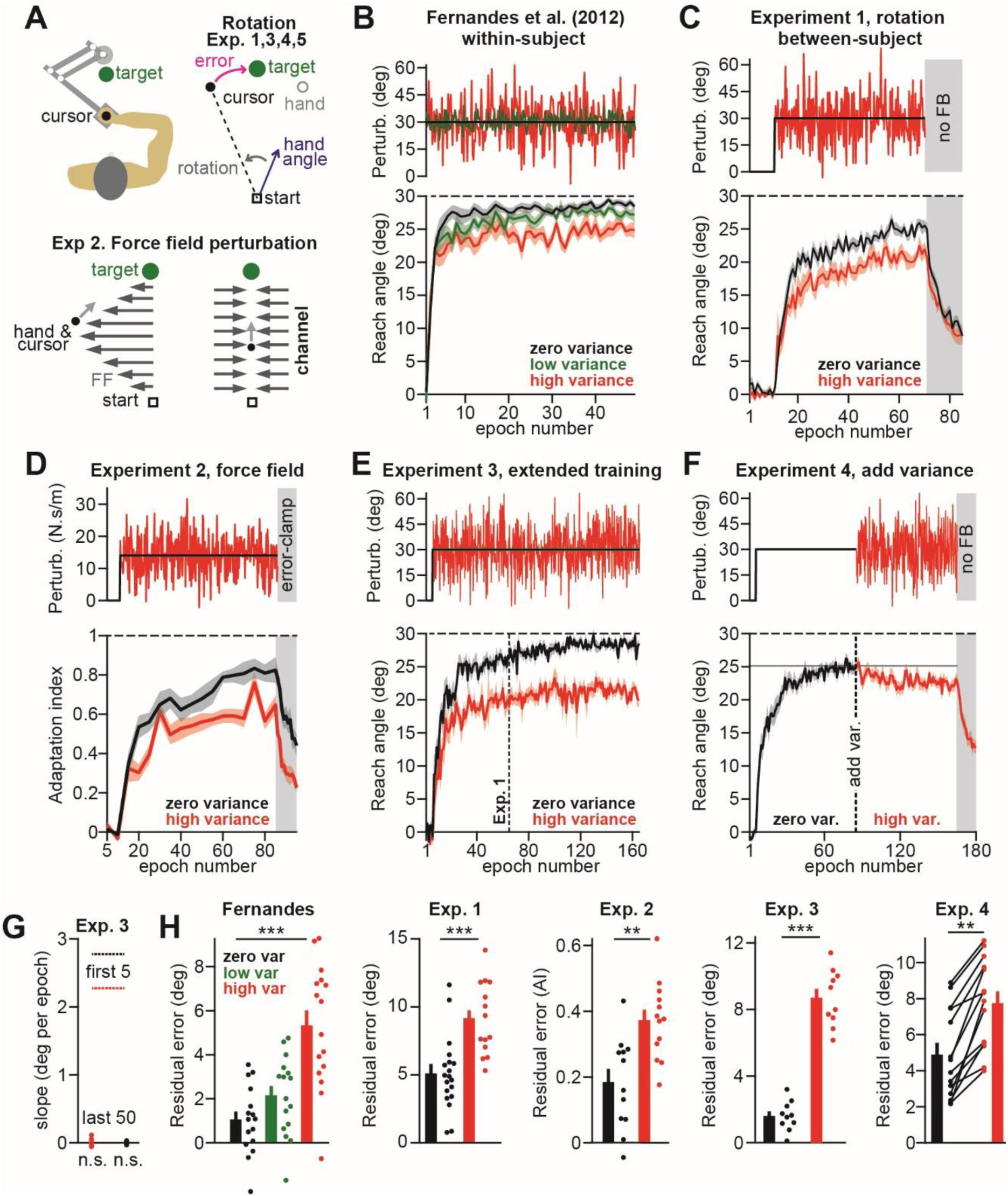
Perturbation variance impairs sensorimotor adaptation. **A**. Schematic of our experiment setup. **B**. Fernandes and colleagues ^19^ measured the reach angle of participants (bottom, n=16) during adaptation to variable visuomotor rotations (top: SD = 0, 4, and 12° for zero, low, and high-variance; mean is 30° for all). Participants demonstrated differing residual errors (reported in inset **H, Fernandes**; median error on the last 48 trials). **C**. In Experiment 1, we repeated the experiment of Fernandes et al. (2012) with a between-subjects design. Participants adapted to a zero (n=19) or high (n=14) variance perturbation (SD = 0 and 12° for zero and high-variance; mean is 30° for both). The residual error is shown in **H, Exp. 1** (median of the last 48 trials). **D**. In Experiment 2, we tested force field adaptation. Occasionally, we measured reaching forces on channel trials that restricted motion of the hand to a straight path. Participants experienced a zero (n=12) or high (n=13) variance perturbation (top: SD = 0 and 6 N-s/m for zero and high-variance; mean = 14 N-s/m for both). We computed an adaptation index on each channel trial (bottom). Residual error (inset **H, Exp. 2**) is one minus mean adaptation index on last 5 error clamp trials. **E**. In Experiment 3, we exposed participants to an extended period of visuomotor rotations (160 epochs = 640 trials). The vertical dashed line indicates the total number of rotation trials in Experiment 1. Participants adapted to a zero (n=19) or high (n=14) variance perturbation (top: SD = 0 and 12° for zero and high-variance; mean is 30° for both). Mean residual error (inset **H, Exp. 3**) was computed over the last 50 epochs. To confirm that performance had reached a plateau, we measured the slope of a line fit to the same period (inset **G**). For comparison, horizontal dashed lines show the mean slope over the first 5 epochs of the perturbation. **F**. In Experiment 4, we adapted participants (n=14) to a zero-variance perturbation, and then abruptly switched to a high-variance perturbation. Residual errors (inset **H, Exp. 4**) were computed over the last 10 epochs of each period. Error bars are mean ± SEM. Statistics denote the result of a repeated-measured ANOVA (**H, Fernandes**) or two- sample t-tests (**H**, all other insets). Statistics: **p<0.01 and ***p<0.001.

To answer this question, we repeated the experiments of Fernandes and colleagues^19^, but with an important difference. In that earlier work, all three perturbation conditions were experienced by the same participants, raising the possibility that prior exposure to the visuomotor rotation could have altered subsequent learning in the other environments^31,34,35^. To avoid this possibility, we recruited different sets of participants for each perturbation condition.

In our experiments, participants held the handle of a robotic arm (Fig. 1A) and reached in a two-dimensional workspace. In Experiment 1, we introduced a visual perturbation and divided the participants into two groups: a zero-variance group (n=19) in which the perturbation magnitude remained invariant at 30° (Fig. 1C, black), and a high-variance group (n=14) in which the perturbation was sampled on each trial from a normal distribution with a mean of 30° and standard deviation of 12° (Fig. 1C, red). Our results confirmed the earlier observation; participants in the zero-variance group learned more than the high-variance group (Fig. 1C, bottom; Fig. 1H, Exp. 1, mean error on last 10 epochs, two-sample t-test, p=0.002; Cohen’s d=1.49).

In Experiment 2, we tested the generality of this observation by measuring how participants responded to variability in force field perturbations (Fig. 1A, Force field). As before, we divided the participants into two groups, a zero-variance group (n=12) in which the perturbation magnitude remained constant at 14 N⋅sec/m (Fig. 1D, top, black), and a high-variance group (n=13) in which the perturbation magnitude was sampled on each trial from a normal distribution with mean 14 N⋅sec/m and standard deviation of 6 N⋅sec/m (Fig. 1D, top, red). To track the learning process, we intermittently measured reach forces during channel trials^36^ (Fig. 1A, channel). As in visuomotor adaptation, variance in the force field perturbation reduced the total amount of learning (Fig. 1D, bottom; Fig. 1H, Exp. 2, mean error on last 5 epochs; two-sample t-test, p=0.001; Cohen’s d=1.46). Thus, perturbation variability altered the extent of adaptation across various modalities of adaptation.

### Perturbation variance limited the total extent of adaptation

An examination of the late stage of training (Figs. 1B-D, bottom) raises the concern that adaptation had not completely saturated; perhaps with additional exposure, adaptation might converge across variance conditions, even eliminating the residual errors. To examine this possibility, we repeated Experiment 1, but this time more than doubled the number of training trials (Fig. 1E). Addition of these trials allowed performance to saturate, as evidenced by the slope of the reach angles (Fig. 1G, slope of the line fit to individual performance over the last 50 epochs was not different than zero; p=0.71 and p=0.83 for the low and high-variance groups). Notably, despite extended training, residual errors persisted (Fig. 1H, Exp. 3, residual errors ± SD on last 50 epochs; zero-variance: 1.7 ± 0.9°; high-variance: 8.7 ± 1.7°; t-test against zero; both groups, p<0.001). We again found that high perturbation variance coincided with an increase in residual error (Fig. 1H, Exp. 3; two-sample t-test, p<0.001; Cohen’s d=5.24).

Did perturbation variability causally alter asymptotic performance? If so, we reasoned that we could switch between two different asymptotic states by changing the perturbation variance mid-experiment. To test this prediction, in Experiment 4 participants (n=14) first adapted to a zero-variance 30° visuomotor perturbation (Fig. 1F, black). With training, performance approached a plateau. We next increased the perturbation variance (while keeping the mean constant) by sampling from a normal distribution with a standard deviation of 12° (Fig. 1F, red). As the perturbation variance increased, reach angles decreased (Fig. 1H, Exp. 4, mean residual error on last 10 epochs; two-sample t-test, p=0.005; Cohen’s d=1.16). Thus, despite having already learned to compensate for much of the perturbation, when perturbation variance increased, residual error increased in every subject (Fig. 1H, Exp. 4).

Together, Experiments 1-4 demonstrated that despite extended practice, motor adaptation suffered from an asymptotic limit, resulting in persistent errors. However, this asymptotic limit was dynamic, responding to the second order statistics of the perturbation.

### Residual errors were a property of the implicit learning system

While reach adaptation can occur despite severe damage to the explicit, conscious learning system of the brain^8^, under normal circumstances performance benefits from both implicit and explicit learning systems ^10,11,37,38^. Therefore, in principle, the residual errors might be due to limitations in implicit learning, explicit learning, or both. To explore this question, we performed a series of experiments that isolated each learning system and measured the effects of perturbation variance on performance.

To isolate the explicit learning system we used a well-documented approach: delayed feedback ^25,26,39,40^. We removed all visual feedback during the movement itself, and only presented the terminal endpoint of the cursor to the participant at a delay of 1 second following movement completion (Experiment 5). This feedback delay is thought to impair implicit learning, at least in part by delaying olivary input to the cerebellar cortex well beyond a plasticity window that peaks at approximately 120 ms^41,42^. As in Experiment 1, we tested participants using perturbations with zero variance and high variance (Fig. 2A). We found that in both groups the increased feedback delay accelerated the learning rate, consistent with the rapid expression of aiming strategies (Fig. 2A, bottom left and right panels). Furthermore, the reaction times greatly increased in all periods of the experiment (Figs. 2A and 2B, bottom). Remarkably, now the subjects compensated perfectly for the mean perturbation and were able to eliminate the residual errors (Fig. 2B, t-test against zero, p=0.512 for zero-variance and p=0.978 for high-variance). Furthermore, in contrast to all prior experiments (Fig. 1), perturbation variability did not have any measurable effects on asymptotic performance (Fig. 2B, bottom). That is, at the end of training, there was no difference in residual error among the zero and high variance groups (Fig. 2B, bar graph, paired t-test, p=0.522). Thus, reach adaptation that putatively relied on explicit learning did not exhibit residual errors, and was not affected by perturbation variance.

**Figure 2.**
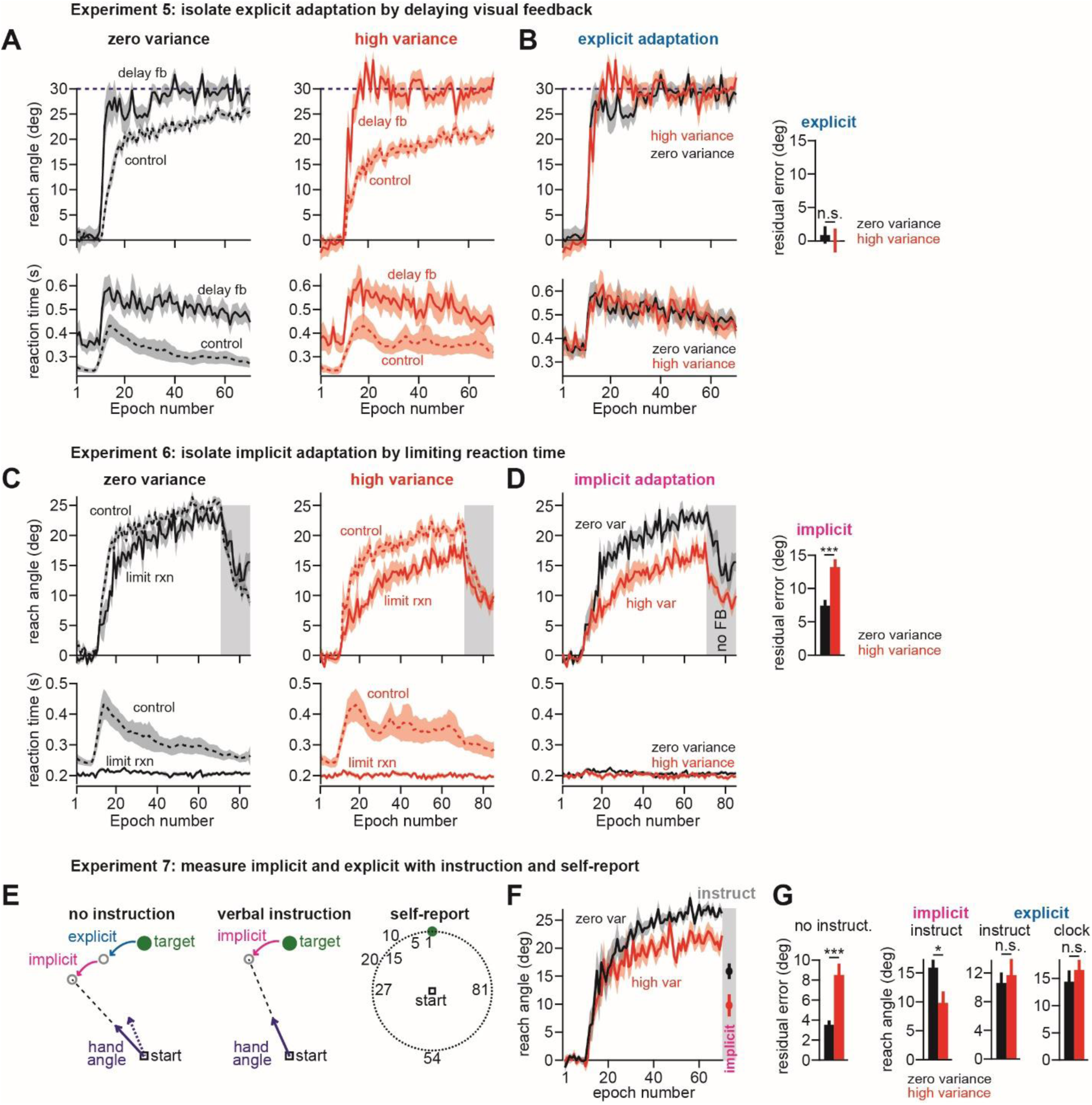
Perturbation variance altered the total extent of implicit, but not explicit adaptation. **A**. We exposed subjects to zero-variance and high-variance visuomotor rotations, under the conditions in Experiment 5 that isolated explicit adaptation (only endpoint feedback, with a delay of approximately 1 second). At left we show the explicit response (solid lines, “delay fb”) to the zero-variance perturbation and at right we show the explicit response to the high-variance perturbation. These responses are compared to the control conditions in Experiment 1. At top we show reach angles and at bottom we show the corresponding reaction times. **B**. Here we compare the explicit response to the zero-variance and high-variance perturbation in Experiment 5. At right, we show the residual error over the last 10 epochs of the rotation period. **C**. We exposed subjects to a zero-variance and high-variance visuomotor rotations, under the conditions in Experiment 6 that isolated implicit adaptation (upper bound on reaction time to prevent the expression of explicit strategies). At left we show the implicit response (solid lines, “limit rxn”) to the zero-variance perturbation and at right we show the implicit response to the high-variance perturbation. These responses are compared to the control conditions in Experiment 1. At top we show reach angles and at bottom we show the corresponding reaction times. **D**. Here we compare the implicit response to the zero-variance and high-variance perturbation in Experiment 6. At right, we show the residual error over the last 10 epochs of the rotation period. **E**. In Experiment 7, we measured the terminal levels of implicit and explicit adaptation after learning using the control conditions in Experiment 1. Normally, learning is composed on implicit and explicit elements (left schematic, no instruction). To isolate the implicit component, we verbally instructed participants to move their hand (not the cursor) through the target without any feedback (middle schematic, verbal instruction). To isolate the explicit component, we asked participants to indicate where they aimed their hand using visual landmarks (right schematic, self-report). **F**. Here we show the reach angle during the learning period in Experiment 7. In the gray region, we show the implicit learning remaining after the verbal instruction. **G**. In column 1, we show the mean residual error in over the last 10 epochs of Experiment 7. In column 2, we show the implicit learning at the end of adaptation that remained after the verbal instruction. In column 3, we obtained the explicit reach angle by subtracting the implicit learning measured after verbal instruction from the total learning curve measured prior to the verbal instruction. In column 4, we show the mean aiming angle self-reported by the participants in each group. Error bars are mean ± SEM. Statistics: *p<0.05, ***p<0.001, and n.s. indicates p>0.05.

This hints that variation in residual errors (Fig. 1) may be due to properties of the implicit learning system. To explore this possibility, we isolated implicit adaptation by severely limiting the time that participants were given to initiate their movement^22,23,43,44^. We did this by imposing a strict upper bound on reaction time and systematically training participants to reach at very low latencies (Experiment 6). As before, we divided participants into two groups: a zero-variance group (n=13) and a high-variance group (n=12) with perturbation statistics identical to that of Experiment 1.

Under normal condition in which there was no constraint on reaction time, introduction of the perturbation led to a dramatic increase in reaction time (Fig. 2C bottom panel, control); participants nearly doubled their preparation time, potentially signaling the expression of explicit strategies. In contrast, in the constrained reaction time group, subjects executed their reach at considerably lower latencies (Fig. 2C bottom panel, limit rxn). In this group, the time required for movement preparation remained roughly constant throughout the experiment, even after the introduction of the perturbation.

As expected, limiting reaction time impaired adaptation. In the zero-variance (Fig. 2C, left panel) and high-variance conditions (Fig. 2C, right panel), performance at short reaction times was stunted relative to control (two-sample t-test on last 10 epochs; p=0.041 and p=0.007 for zero and high-variance; Cohen’s d=0.77 and 1.17 for zero and high-variance), consistent with the removal of explicit aiming strategies. Critically, the residual errors expressed by the isolated implicit system were clearly affected by increased perturbation variance (Fig. 2D); the total extent of learning was reduced by approximately 5° (Fig. 2D, bar graph, difference in residual errors during the last 10 epochs, two-sample t-test, p<0.001, Cohen’s d=1.53). Thus, whereas perturbation variance did not affect the explicit system, it severely impaired the implicit system.

Under normal circumstances both the implicit and explicit systems contribute to adaptation. This is illustrated schematically in Fig. 2E (left subplot); when a target is presented, explicit-based learning rotates the target by some amount, and the implicit system provides a subconscious recalibration, resulting in the eventual reach angle. Our results in Fig. 2B suggest that perturbation variance does not affect the explicit system. If this is true, then during normal adaptation in which both implicit and explicit systems contribute to learning, assay of implicit and explicit contributions should show that perturbation variance impairs only the implicit component, not explicit strategy.

We tested this prediction by performing a control experiment (Experiment 7). Participants were divided into two groups (zero-variance, n=9, and high-variance, n=9) and experienced perturbations with statistics matching those of Experiment 1. As expected, the addition of perturbation variance reduced the total extent of adaptation (Fig. 2F, also shown in Fig. 2G, no instruct mean residual error over last 10 epochs; two-sample t-test, p<0.001, Cohen’s d=1.91). To determine if these differences in performance were caused by the effect of perturbation variance on the implicit system, explicit system, or both, we conducted two assays at the end of the training period. First, we verbally instructed participants that the cursor would be removed on the next several trials, and their goal was to move their hand straight through the target, without trying to compensate for any rotation that they had experienced (Fig. 2E, verbal instruction). Such an instruction eliminates explicit aiming, thus isolating the amount of implicit adaptation^12,13^ (Fig. 2F, gray region). By subtracting this implicit angle from the reach angle measured prior to the verbal instruction, we also estimated the extent to which participants were explicitly aiming their reach angle at the end of the adaptation period. We found that in the high-variance condition, the implicit system had learned less than in the zero-variance condition (Fig. 2G, implicit instruct, two-sample t-test, p=0.023, Cohen’s d=1.19). In contrast, explicit aiming was unaltered by perturbation variance (Fig. 2G, explicit, instruct, two-sample t-test, p=0.69), thus confirming the results in Fig. 2B. That is, perturbation variance appeared to impair only the implicit system.

Next, we followed the implicit probe with another assay to measure explicit aiming. Participants were shown a target as well as a ring of small dots each labeled with an alphanumeric string (Fig. 2E, self-report). At the end of the perturbation period we asked them to report the angle toward which they aimed their hand (using the small dots as a guide). We again found that perturbation variance had no effect on explicit aiming (Fig. 2G, explicit clock, two-sample t-test, p=0.45).

In summary, when learning relied mainly on the explicit system, performance did not suffer from residual errors (Fig. 2A), and was unaffected by perturbation variability. In contrast, when learning relied mainly on the implicit system, performance exhibited residual errors, and was strongly affected by perturbation variability (Fig. 2C). When the two-learning system operated together, perturbation variance affected only the implicit system (Fig. 2G). Thus, change in residual error appeared to be caused by properties of the implicit system, properties that were sensitive to the second order statistics of the perturbation.

### Perturbation variance reduced error sensitivity, but not forgetting rates

Why does the implicit system exhibit an inability to completely eliminate performance errors, and why is this impairment exacerbated by perturbation variance? In principle, steady-state errors arise because performance is driven by an interaction between two opposing forces, error-based learning, and trial-to-trial forgetting^17,27–30^ (Fig. 3A). In this model, performance saturates because as training progresses, errors which drive the learning process become small enough that there is a balance between forgetting and learning (see *Methods*). At this stage learning appears to stop, even though residual errors remain. Perturbation variance might have affected forgetting rates, or error sensitivity (Fig. 3B and 3C). The implicit system learns with error sensitivity *b*_*i*_, and exhibits trial-to-trial retention specified by *a*_*i*_. Similarly, the explicit system learns with error sensitivity *b*_*e*_, and exhibits trial-to-trial retention specified by *a*_*e*_. Does perturbation variance affect error sensitivity, forgetting, or both?

**Figure 3.**
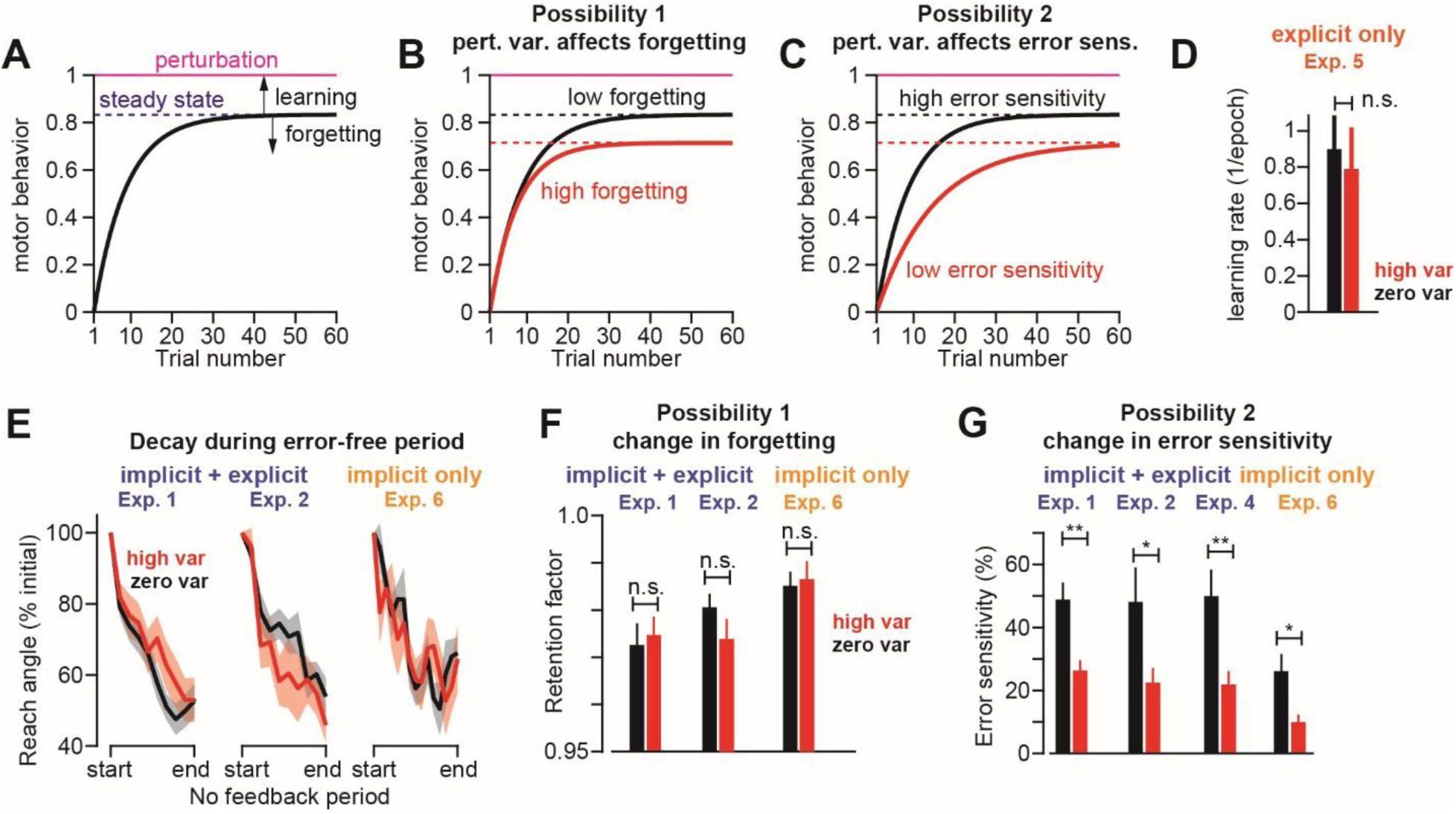
Perturbation variance decreases error sensitivity, not decay rates. **A**. State-space model of adaptation predicts that learning will reach an asymptote when the amount of learning from an error exactly counterbalances the amount of forgetting that occurs between trials. The plot demonstrates the behavior of such a model during adaptation to a perturbation of unit 1. According to the model, changes in asymptotic levels of performance can occur because of changes in forgetting (**B**, Possibility 1 schematic; *a* = 0.98 for low forgetting and 0.96 for high forgetting), or changes in error sensitivity (**C**, Possibility 2 schematic; *b* = 0.05 for low error sensitivity and 0.1 for high error sensitivity). **D**. Perturbation variance had no effect on the rate of explicit learning. Here we show the learning rate in the zero-variance and high-variance groups of Experiment 5 quantified with an exponential fit to individual participant behavior **E**. To test Possibility 1, we measured the retention during error-free periods at the end of Experiments 1 (left), 2 (middle), and 6 (right). We normalized reach angle to the first trial in the no-feedback period. Each point on the x-axis is a cycle of 4 trials. **F**. We measured the retention factor during error-free periods in each experiment depicted in **E**. We found no difference in retention for the zero-variance and high-variance groups. **G**. To test Possibility 2, we measured sensitivity to error in each experiment that terminated with an error- free period. Error sensitivity was greater for the zero-variance perturbation in every experiment. Error bars are mean ± SEM. Statistics: *p<0.05, **p<0.01, and n.s. indicates no statistical significance.

First, we consider explicit adaptation. Because explicit strategies did not exhibit residual errors in either the zero-variance and high-variance environments (Fig. 2B), we can infer that the explicit system does not suffer from trial-to-trial forgetting. In the state-space framework, this implies that *a*_*e*_ ≈ 1 irrespective of perturbation variability (see *Methods*). Did perturbation variance affect error sensitivity of the explicit system? To answer this question, we examined the data in Exp. 5 and found that the learning rate was not different among the groups that learned with zero or high perturbation variance (Fig. 3D, paired t-test, p=0.715). These results suggest that for the explicit system, both trial-to-trial forgetting and error sensitivity are unaltered by perturbation variance.

We next focused on the implicit system and began by estimating the forgetting rate of each participant in the error-free movement period at the end of each experiment (gray region in Figs. 1C, 1D, 1F). During these periods, behavior naturally decayed towards the baseline (Fig. 3E), thus providing a way to isolate the rate of trial-by-trial forgetting (i.e., the rate of decay of behavior). Interestingly, we found that in all experiments, the rate of forgetting was unchanged by perturbation variability (Fig. 3E, two-sample t-test; Exp. 1, p=0.72; Exp. 2, p=0.19; Exp. 6, p=0.79). Critically, when we isolated the implicit system, the rate of forgetting was unaffected by perturbation variance (Exp. 6, Figs. 3E and 3F). Therefore, perturbation variance did not affect the forgetting rate in the implicit system.

Next, we empirically estimated error sensitivity in the various experiments. To do this, we calculated the difference between the reach angle in pairs of consecutive trials (adjusting for forgetting) and divided this by the error experienced on the first of the two trials. By definition, this quotient represents one’s sensitivity to error, i.e., the fraction of the error that is compensated for on the next trial. In sharp contrast to forgetting rates, we found consistent differences in error sensitivity between the zero and high perturbation variance groups; in all experiments, participants in the zero-variance groups exhibited an error sensitivity nearly twice that of individuals in the high-variance groups (Fig. 3G: two-sample t-test; Exp. 1, p=0.002, Cohen’s d=1.18; Exp. 2, p=0.039, Cohen’s d=0.87; Exp. 4, p=0.006, Cohen’s d=1.12; Exp. 6, p=0.016, Cohen’s d=1.05). Importantly, when we isolated the implicit system, error sensitivity was significantly reduced by variance (Fig. 3G).

In summary, perturbation variance had no effect on the explicit system, but reduced error sensitivity of the implicit system. This suggests that residual errors increased with high-variance perturbations because the variance somehow reduced the error sensitivity of the implicit system.

### Perturbation variance reduced the ability to learn from small errors, not large errors

Our quantification of error sensitivity in Fig. 3G made the assumption that the brain is equally sensitive to errors of all sizes. However, it is well-documented that error sensitivity varies with the magnitude of error; one tends to learn proportionally more from small errors^12,13,46,47^. In other words, error sensitivity is not constant, but declines as error size increases. How did perturbation variance alter the functional relationship between error magnitude and sensitivity to error?

To answer this question, we re-estimated error sensitivity, but this time controlled for the magnitude of error. We placed pairs of consecutive movements into bins according to the error experienced on the first trial, and then calculated error sensitivity within each bin. As expected, in both zero-variance and high-variance conditions, as error size increased, error sensitivity decreased (Fig. 4A, left; mixed-ANOVA, within-subjects effect of error size, F=22.1, p<0.001, 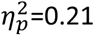). This confirmed that indeed, people tended to learn proportionally less from larger errors. However, for a given error size, the high-variance perturbation group exhibited lower error sensitivity than the zero-variance group (Fig. 4A, left; mixed-ANOVA, between-subjects effect of perturbation variance, F=14.7, p<0.001, 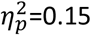).

**Figure 4.**
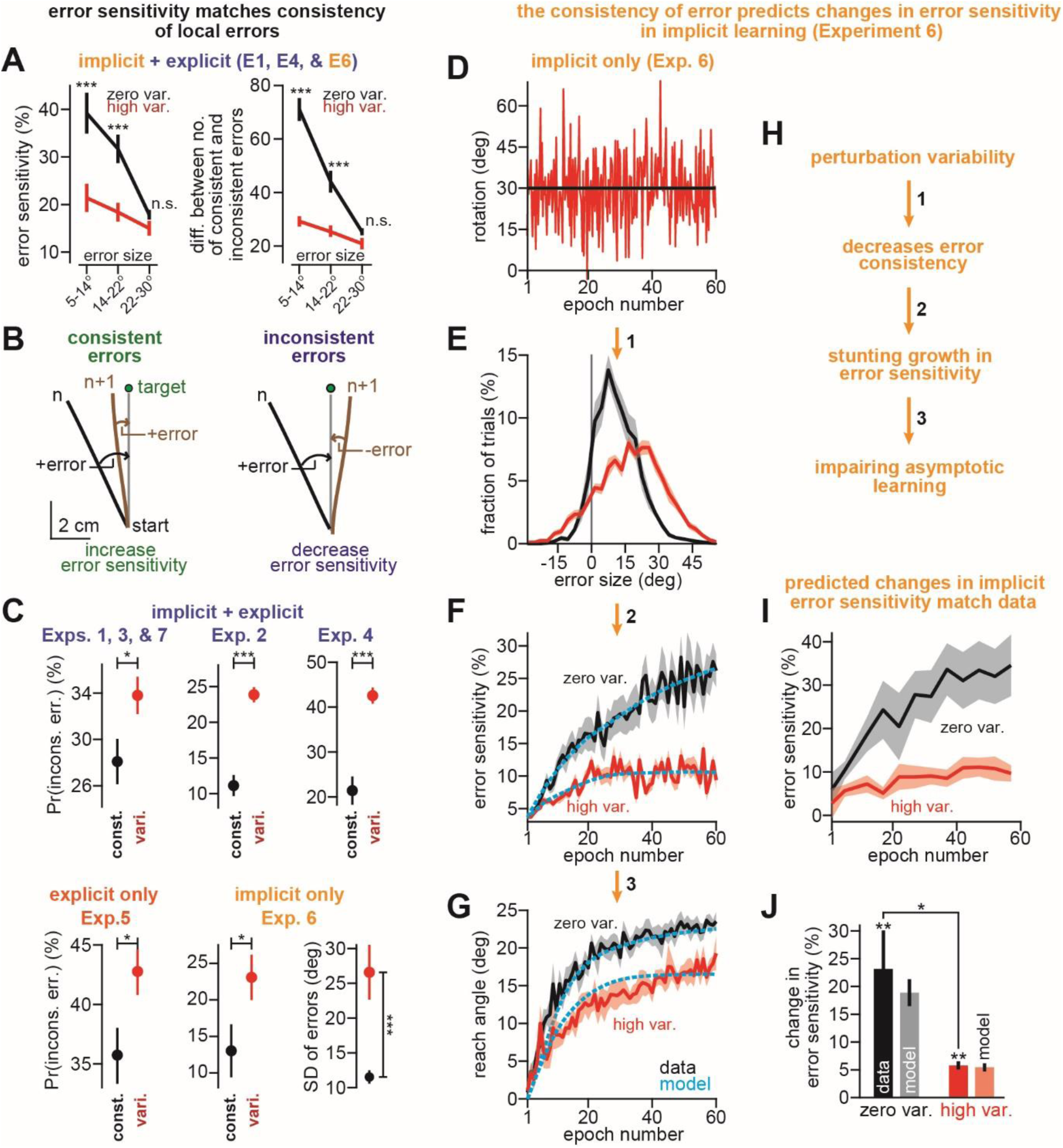
Spatiotemporal variation in error sensitivity is predicted by the consistency of error. **A**. Left: to determine how error sensitivity varied as a function of error size, we sorted pairs of movements into different bins according to the size of the error on the first movement. Next, we computed the mean error sensitivity across all trials within each error size bin. To increase power, we combined participants across all visuomotor rotation experiments with an implicit learning component and an error-free period in which retention could be independently measured (Experiments 1, 4, and 6). Right: the difference between the number of consistent and inconsistent errors during adaptation to the visuomotor rotation for the error sensitivity measurements at left. **B**. We considered the possibility that the trial-to-trial consistency of errors caused changes in error sensitivity. Consistent errors (left) are consecutive pairs of trials where the errors have the same sign. Inconsistent errors (right) are consecutive pairs of trials where the errors have opposite signs. The black and brown traces show example reach trajectories from a single participant. **C**. We measured the total fraction of inconsistent error trials. The high-variance perturbation caused a higher probability of inconsistent errors in every experiment. Each inset shows the probability of experience in inconsistent error for a given experiment, or set of experiments. The only exception is at bottom- right. Here we show the standard deviations of the error distributions corresponding to the zero-variance and high-variance groups in Experiment 6 (distributions shown in **E**). **D-G**. Here we break down the behavior of our decaying memory of errors model for the implicit-only behavior recorded in Experiment 6. Addition of variability to the perturbation (**D**) altered the distribution of errors experienced in the high-variance group (**E**). Using the error sequences (summarized in **E**) we used Eq. (1) to predict how implicit error sensitivity should vary as a function of trial. The mean error sensitivity timecourse predicted by the model is shown in **F**. The noisy solid curves show the mean timecourse across participants. The dashed blue lines show a smoothed version used for simulation of behavior. In **G**, we simulate the implicit learning curves predicted by Eq. (1) using the implicit error sensitivity depicted in **F** and the state-space model in Eqs. (3-5). **H**. Here we provide a verbal schematic depicting how the decaying memory of errors model (Eq. (1)) translates changes in perturbation variance to differences in error sensitivity, and ultimately, to two different asymptotic states of learning. **I**. Here we show the timecourse of error sensitivity empirically measured across participants in Experiment 6. **J**. Here we show the change in error sensitivity measured from the start to the end of learning for the measured behavior depicted in **I** (left bars in the zero var. and high var. groups) and that predicted by Eq. (1) depicted in **F** (right bars in the zero var. and high var. groups). Error bars are mean ± SEM. For **A**, we used a mixed-ANOVA followed by post-hoc two-sample t-tests with Bonferroni corrections. In **C** and **J**, two-sample or paired t-tests were used for statistical testing. Statistics: *p<0.05, **p<0.01, ***p<0.001 and n.s. indicates no statistical significance.

This analysis revealed an interesting pattern; increased perturbation variance reduced the ability to learn from small errors (<20°), but had no effect on the ability to learn from larger errors (>20°) (Fig. 4A left; post-hoc testing with t-test adjusted with Bonferroni correction, p<0.001 and Cohen’s d=0.72 for 5-14°, p<0.001 and Cohen’s d=0.79 for 14-22°, and p=0.53 for 12-30°). Why should increases in perturbation variance selectively affect learning from smaller errors, but not larger errors?

### The spatial pattern of error sensitivity follows the consistency of error

A model of sensorimotor adaptation^31^ posits that the brain adjusts its sensitivity to error in response to the consistency of past errors. In this memory of errors model, when the error on trial *n* has the same sign as the error on trial *n*+1, it signals that the brain has undercompensated for error on trial *n*, and so should increase sensitivity to that error (Fig. 4B, left). Conversely, when the errors in two consecutive trials differ in sign, the brain has overcompensated for the first error, and so should decrease sensitivity to that error (Fig. 4B, right). These changes in error sensitivity occur locally, meaning that the brain can simultaneously increase sensitivity to one error size, while decreasing sensitivity to another^31^. Thus, in the context of a variable perturbation, the memory of errors model provides an interesting prediction; perturbation variance alters the consistency of errors, producing less consistency for some error sizes (smaller ones) but not others (larger ones).

We tested this idea by quantifying consistency of error as a function of its size. Indeed, we found that in the high-variance group there was a higher probability of experiencing an inconsistent error (Figs. 4C; Exps. 1, 3, & 7, p=0.029, Cohen’s d=0.53; Exp. 2, p<0.001, Cohen’s d=2.84; Exp. 4, p<0.001, Cohen’s d=2.22; Exp. 5, p=0.031, d=0.90; Exp. 6, p=0.048, Cohen’s d=0.84). Moreover, when we binned the data based on error size, the differences in the relative number of consistent and inconsistent errors exhibited a striking pattern that mirrored error sensitivity patterns (Fig. 4A, right; mixed-ANOVA, between-subjects effect of perturbation variance, F=60.5, p<0.001, 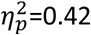; within-subjects effect of error size, F=54.4, p<0.001, 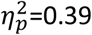). For smaller errors, the zero-variance group had more consistent error events and fewer inconsistent error events than the high-variance group (Fig. 4A, right; post-hoc testing with t-test adjusted with Bonferroni correction, p<0.001 and Cohen’s d=1.83 for 5-14°, p<0.001 and Cohen’s d=0.85 for 14-22°). However, for large errors, there was no difference in the relative consistency (Fig. 4A, right; post-hoc testing with t-test adjusted with Bonferroni correction, p=0.16).

In summary, as perturbation variance increased, there was a reduction in the trial-to-trial consistency of small errors, but not large errors (Fig. 4A, right). Coincident with these changes in the history of errors, there was a reduction in the error sensitivity for small errors, but not large errors (Fig. 4A, left). These results raised the possibility that changes in error sensitivity in the implicit system (Fig. 3G) were due to the history of errors that each participant had experienced throughout training. To explore this question, we further analyzed the data in the framework of the memory of errors model.

### The temporal pattern of error sensitivity follows the consistency of error

The memory of errors model^31^ posits that error sensitivity changes during training as a function of the specific sequence of errors that each participant has experienced:

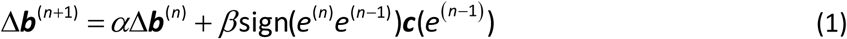

Here, **Δ*b*** is a vector whose elements represent the change in error sensitivity within different patches of the error space (5° bins spaced evenly between errors of -100° and 100°). Eq. (1) describes changes in error sensitivity in terms of two forces: learning and decay. Learning is encapsulated in the right-most term, which increases error sensitivity when consecutive errors are consistent. The rate of this increase is determined by the parameter *β*. Error sensitivity increases only for error sizes close to the error experienced on the first of the two consecutive trials (controlled by the vector ***c***, see *Methods*). Decay is encapsulated by the parameter α, which like the retention factor (Fig. 3F), determines how strongly the memory of past errors is retained from one trial to the next.

We focused on Exp. 6 where we isolated implicit learning. For each participant, we used their actual sequence of errors to predict how error sensitivity should vary for a given error size throughout training. When variability was added to the perturbation (Fig. 4D), this changed the statistics of error (Fig. 4E, Step 1 in Fig. 4H). The error distribution widened (i.e., became more variable; Fig. 4C, Exp. 6, SD of errors, two-sample t-test, p<0.001, Cohen’s d=1.55), and also exhibited an increased probability of experiencing inconsistent errors (Fig. 4C, Exp. 6, left, two-sample t-test, p=0.048, Cohen’s d=0.84). Because of these changes in the underlying error distribution, Eq. (1) predicted that implicit error sensitivity should diverge over time in the zero-variance and high-variance environments (blue curves in Fig. 4F, Step 2 in Fig. 4H). Finally, because in the high-variance group the implicit error sensitivity saturated prematurely, the process of error-based learning was suppressed, thereby reducing the total extent of implicit adaptation (Fig. 4G, Step 3 in Fig. 4H). The cascade of these processes predicted behavior that closely matched the observed reach angles (Fig. 4G).

The model made the unexpected prediction that error sensitivity should increase during training, but at a slower rate for the high-variance group (Fig. 4F, blue curves). Despite high perturbation variance, the experience of consistent errors (Fig. 4B, left) remained more probable than inconsistent errors (Fig. 4C, Exp. 6, left). Therefore, Eq. (1) made the surprising prediction that error sensitivity should increase in both the zero-variance and the high-variance environments, but less so in the high-variance case (Fig. 4F; Fig. 4J, model).

To test for this, we empirically calculated implicit error sensitivity as a function of trial in the zero-variance and the high-variance groups. Critically, we found that implicit error sensitivity started at similar levels in the zero-variance and high-variance environments (two-sample t-test on error sensitivity over first 10 epochs, p=0.20), but diverged over time (Fig. 4I). Both of these predictions matched the observed implicit time courses (Fig. 4J, data). Implicit error sensitivity increased during exposure to the zero-variance perturbation (Fig. 4J, zero var., left bar; paired t-test, p=0.006, Cohen’s d=0.93), and during the high-variance perturbation (Fig. 4J, high var., left bar; paired t-test, p<0.001, Cohen’s d=2.21). However, the growth rate was stunted in the high-variance group relative to the zero-variance group (Fig. 4J, compare left bars in zero var. and high var.; two-sample t-test, p=0.025, Cohen’s d=0.96).

In summary, our model predicted that implicit error sensitivity should increase in response to the more consistent history of errors in the zero-variance perturbation condition. It also predicted that introducing variance into the perturbation should not decrease implicit error sensitivity, but rather stunt its growth. Our measurements confirmed both of these predictions. Thus, the implicit process of adaptation behaved in a manner consistent with an error sensitivity that depended on a decaying history of past errors.

### Generality of the model and its predictions

Our model makes the general prediction that the specific sequence of errors that the subject experiences affects the error sensitivity of the implicit system, which in turn produces an inability to eliminate residual errors. To test the generality of this prediction, we considered two important data sets in another implicit learning paradigm: non-zero error-clamp condition and dissolution of savings during saccade adaptation.

Robinson and colleagues^6^ adapted monkeys to a saccadic perturbation in which the error on every trial was fixed to -1° independent of the monkey’s motor output (Fig. 5A, top). Critically, despite the fact that error never changed, performance nevertheless reached a saturation point (Fig. 5A, middle). We simulated Eq. (1) and found a similar behavior: despite complete error consistency, the presence of decay (*α*<1) caused error sensitivity to saturate over time (Fig. 5A, bottom). Because error sensitivity saturated, so too did behavior (Fig. 5A, middle). In contrast, if decay was not present (*α*=1), error-sensitivity grew unbounded, and model predictions did not match the data. Therefore, the memory of errors model exhibits saturation in performance during non-zero error-clamp conditions, but only if there is decay in the memory of errors.

**Figure 5.**
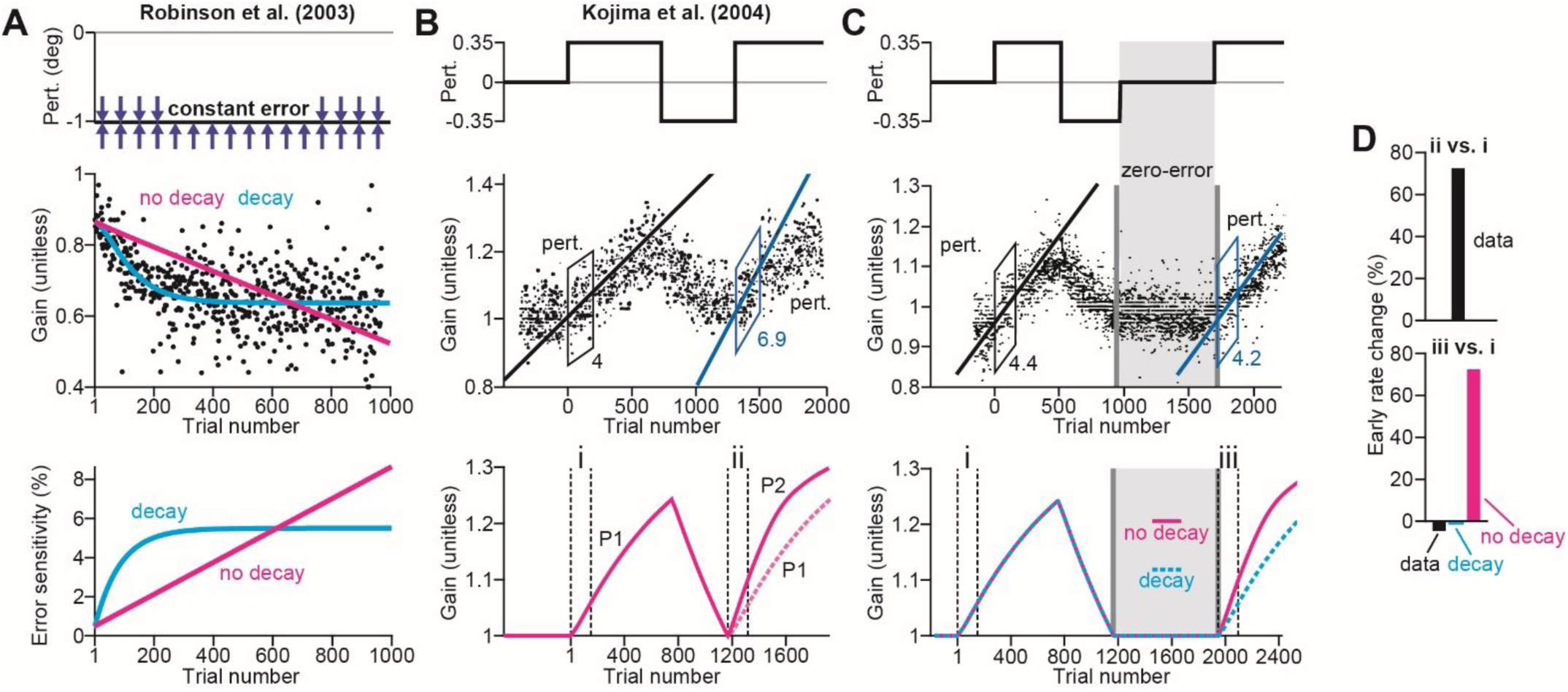
The memory of errors decays over time. **A**. Data were obtained from Robinson and colleagues^6^. Monkeys were adapted to a gain-down saccade perturbation. The error on each trial was fixed to -1° as shown at top. The black points in the middle inset the saccadic gain recorded on each trial. We fit the “decay” and “no decay” models to the trial-by-trial behavior in the least-squares sense. The decay model fit is shown in blue. The no decay model fit is shown in magenta. At bottom, we show the timecourse of error sensitivity predicted by the decay and no decay models. **B**. Data were obtained from Kojima and colleagues^32^. The authors adapted monkeys to a gain-up perturbation, followed by a gain-down perturbation, followed by a re-exposure to the gain-up perturbation. Paradigm is shown at top. Saccadic gain recorded on each trial during a representative session is shown at middle. The black and blue regression lines represent the linear fit to the first 150 trials during the initial exposure and re-exposure to the gain-up perturbation. Behavior exhibited savings in this paradigm, as indicated by the slope of the regression lines. At bottom, we show the output of the no decay memory of errors model described by Herzfeld and colleagues^31^. P1 refers to the first gain-up perturbation. P2 refers to the second gain-up perturbation. **C**. Data were obtained from Kojima and colleagues^32^. In a second experiment, monkeys adapted to a similar perturbation schedule as in **A**, only this time a long period of zero perturbation trials was added prior to the second gain-up adaptation period (shown at top; zero-error). Trial-by-trial saccadic gain is shown at middle. The regression lines indicate the slope of a linear fit to the first 150 trials during the initial exposure and re-exposure to the gain-up perturbation. Note that the “zero-error” period led to the loss of savings as indicated by the slope of the regression lines. At bottom, we show the behavior predicted by the “no decay” model where no decay in error sensitivity is permitted over the zero-error period (solid magenta line). In addition, we simulated a “decay” model, in which error sensitivity decayed during the zero-error period. **D**. We quantified the slope of adaptation in **C** by fitting a line to the behavior of the “no decay” and “decay” models over the periods labeled “i” and “iii”. At top, we show the percent change in slope from “i” to “ii” present in the actual data. At bottom, we show the percent change in slope from “i” to “iii” present in the actual data, the “decay” model, and the “no decay” model.

Finally, we considered a classic experiment that demonstrated savings, but only if the block of re-exposure was temporally close to the block of original exposure, and not if the two were separated by a long washout period. Kojima and colleagues^32^ exposed monkeys to a 3.5° visual perturbation, then a -3.5° perturbation, followed by re-exposure to the original 3.5° perturbation (Fig. 5B, no zero-error period, top). This paradigm elicited savings, i.e., a faster rate of re-learning ^29,31,48,49^ (Fig. 5B, middle; compare initial rates of learning denoted by the linear fits). However, when a long period of washout (no perturbation trials), savings was abolished (Fig. 5C, middle; compare initial rates of learning denoted by the linear fits). We simulated the behavior predicted by Eq. (1) and found that when the number of trials between initial exposure and re-exposure was short (Fig. 5B), the model predicted increased error sensitivity during the re-exposure period, correctly producing savings (Fig. 5B, bottom; compare P1 and P2). However, when the temporal distance was long, the model now predicted no savings upon re-exposure, but only when the memory of errors experienced decay. Therefore, to account for the dissolution of savings, the memory of errors model must decay over time.

In summary, Eq. (1) captures an important duality between the adaptation of behavior and the adaptation of error sensitivity; the dynamics of each are controlled by a competition between learning and forgetting. A decaying memory of errors model not only accounted for the implicit response to variable perturbations (Fig. 4), but also the saturation of learning observed in error-clamp^4^, and the dissolution of savings over long error-free periods^5,9^.

## Discussion

Across numerous paradigms, adaptation exhibits a consistent property; even after prolonged training, learning appears to stop, leaving behind residual errors^2,4,5,7^. Curiously, residual errors depend on the second order statistics of the perturbation; perturbation variance increases residual errors, seemingly impairing adaptation. Here, we find that residual errors are a feature of behavior that arises from the implicit learning system; when reach adaptation depends solely on the explicit system, behavior does not exhibit residual errors. The reason why the implicit system exhibits residual errors is because its error sensitivity depends on the history of errors that the subject has experienced. When perturbation variance is low, errors are temporally consistent, resulting in an increase in the error sensitivity of the implicit system. When perturbation variance is high, errors are temporally inconsistent, causing error sensitivity to rise more slowly in the implicit system. Eventually these up-regulations in error sensitivity strike an equilibrium with forgetting, causing performance to saturate and produce residual errors that cannot be eliminated. Thus, residual errors are a limitation of behavior that arises because the implicit learning system has an error sensitivity that varies with the history of past errors.

### A memory of errors in implicit learning

Motor adaptation is supported by both implicit and explicit systems^8,10,11,37,38^. With the exception of one report^50^, most if not all previous studies^43,51–55^ have suggested that the implicit system is inflexible, has a response to error that does not change with training, and saturates at levels that are identical across perturbations^56^ or error sizes^12,13^.

Our results alter these prevailing views. Using various techniques such as direct verbal instruction^12,13^, reports of explicit aiming angles^9^, limiting movement preparation time^22,23,43,44^, and delaying visual feedback^25,26,39,40^, we found substantial evidence that implicit learning is flexible; its error sensitivity is modulated by the history of past errors. Specifically, both our model and empirical measurements demonstrated that implicit error sensitivity tends to increase with exposure to a perturbation, even when this perturbation is highly variable. In other words, the effect of perturbation variability is not to reduce error sensitivity, but to limit its potential growth. We expect that under natural circumstances, variability in disturbances, the production of a movement, and the process of learning from error^57^, all contribute to the amount of change in implicit error sensitivity and thus the asymptotic behavior of implicit learning.

### Asymptotic behavior of explicit strategies

In contrast to implicit learning, we found that perturbation variability had no effect on explicit strategy, neither decreasing the explicit rate of learning (Fig. 3D), nor its asymptotic performance (Fig. 2D). With that said, many other studies have documented considerable flexibility in the expression of explicit strategy. For example, explicit processes are known to strongly contribute to savings, at least in the context of visuomotor rotation^43,52^. In addition, age-related declines in explicit learning^14,15^ lead to deficits in the total extent of adaptation^16^. Furthermore, manipulations to visual feedback recruit explicit reinforcement learning mechanisms that modulate asymptotic behavior^17^.

Here we found that adding variability to a visual perturbation did not alter the dynamics of explicit learning. However, a recent report^54^ demonstrated that when environmental consistency is added via a random walk, the rate of explicit learning, but not implicit learning, increases. Apparent discrepancies between these observations may relate to methodological differences. In the earlier report, endpoint feedback was provided at delays ranging from 600-2500ms, whereas in our work, participants were provided continuous feedback of the cursor with no added delay (excepting Exp. 5). Therefore, we might expect that this earlier report used conditions that more strongly engaged explicit systems and hindered implicit learning^26,40–42^.

In general, methodological differences across the literature^45^ make it challenging to understand the context-dependent nature by which explicit strategies contribute to asymptotic performance. In some cases, explicit learning reaches a peak early in training, and then declines with further training^9^, reminiscent of a learning system with incomplete retention. Here we found that the explicit system is capable of complete elimination of residual errors, exhibiting no trial-based forgetting (Exp. 5). Even though explicit systems are capable of completely compensating for error (Exp. 5), under normal conditions they do not do so (Exps. 1-4,7). This observation might partially be explained by a recent report^18^ which demonstrates that explicit systems can eliminate residual errors when preparation time is prolonged. Such an interpretation would be consistent with the increased reaction times exhibited in Exp. 5.

In summary, our data support the inclusive view that both implicit and explicit processes change their response to error, and together determine the total extent of sensorimotor adaptation.

### Alternate mechanisms that limit implicit adaptation

Curiously, our conclusions run somewhat counter to recent reports that have engaged implicit systems with invariant target-cursor errors (i.e., constant error-clamp)^12,13^. While we observed fluidity in the extent of implicit adaptation, implicit processes appear to reach similar asymptotic levels when they are driven by fixed errors between the target and cursor. It may be that the total extent of implicit adaptation is limited by an external ceiling in correction that is reached when errors are completely consistent from one trial to the next. But in traditional cases where the consistency of error (Fig. 4B) decreases as adaptation nears its asymptote (thus halting increases in error sensitivity), mechanisms of decay and error-based learning together control the terminal amount of implicit learning.

With that said, it should be noted that there are fundamental differences in the error signals that drive learning in the traditional rotation paradigm used in this study, versus those that employ an invariant error-clamp condition. Implicit systems appear to learn from both hand-cursor error^37^, as well as target-cursor error^24^. In traditional rotation paradigms, the hand-cursor error is constant over time, but the target-cursor error (i.e., task error) decreases over time. In constant error-clamp paradigms^12,13^, the hand-cursor error increases over time, but the target-cursor error remains constant over time. Given these fundamental differences in error signals, it is possible that different rotation paradigms engage different implicit systems. For example, back-of-the-envelope calculations (see *Methods*) indicate that the asymptotic level of implicit learning measured in the constant-clamp paradigm is considerably more variable than that measured under reaction time restrictions in Experiment 6; the standard deviation across participants was ∼300% greater at asymptote for constant-error clamp^13^ versus the limited reaction time condition. It seems unlikely that implicit recalibrations are driven by the same system across each of these tasks.

Finally, it may be that proprioception plays a role in limiting the extent of implicit recalibration, as noted in these earlier studies^13^. In the constant error-clamp condition, the proprioceptive mismatch between the cursor location and the hand position increases as the participant adapts to the perturbation. In traditional adaptation paradigms, this proprioceptive error is fixed. It may be the case that these proprioceptive signals play a modulatory role in limiting the total amount of implicit recalibration in the context of visuomotor adaptation.

### The extent of adaptation is altered by a decaying memory of errors

While many studies have shown that one’s rate of learning^58,59^ can be altered in numerous contexts such as savings^7,29,31,32,34,48,49^, meta-learning^31,48^, and anterograde interference^60^, we know comparatively little about how the brain controls the total amount of adaptation achieved with prolonged exposures to a perturbation. State-space models of learning^27,61^ predict that asymptotic levels of performance are set by the balancing of learning and forgetting^17,62^. Here, we found evidence that error sensitivity is also maintained by the same two processes (Fig. 5).

In our model (Eq. 1) error sensitivity exhibits both consistency-driven modulation as well as trial-based decay. That is, a memory of past errors is both acquired and forgotten over time. The original model^31^ only considered the process of acquisition, not decay. Without decay, it is not possible to account for residual errors. Decay of error sensitivity is evident in the learned response to constant error-clamp conditions^6,12,13,64,65^. When subjects are exposed to the same error time and time again, the decay-free memory of errors model increases error sensitivity without bound (Fig. 5B). Instead, a decaying memory of errors reaches saturation in error sensitivity, and thus produces residual errors.

Consider also the fact that the total extent of learning is often similar during the first and second exposures to a perturbation, even though learning is faster during the second exposure^29,33,66^. Why are residual errors equal, if error sensitivity is higher upon re-exposure? Eq. (1) offers an explanation; while increased error sensitivity leads to faster initial learning upon re-exposure, if error sensitivity decays at the same rate during each exposure, error sensitivity will reach the same steady-state level irrespective of its initial magnitude.

Perhaps the most direct evidence for error sensitivity decay is the loss of savings after long periods of washout (Fig. 5C). That is, adaptation is faster with re-exposure to a perturbation (Fig. 5B), but not when perturbations are separated by long periods of washout^32^. Our model suggests that this dissolution of savings^32,33^ is caused by gradual decay in error sensitivity over error-free periods. While not explored here, we speculate that decay in error sensitivity is more rapid after a movement, than with the passage of time alone. For example, with time alone memory decays, but the rate of re- learning remains elevated^32,33,63^, even after long breaks on the order of a day^55,63^.

### Alternate models

Perturbation variance could also affect uncertainty of the learner. Over the past two decades, numerous studies^28,58,67^ have used a Kalman filter^68^ to study the relationship between uncertainty and learning rate. The Kalman filter describes the optimal way in which an observer should adjust their rate of learning in response to different sources of variability. This Bayesian framework has proved useful in understanding the slowing of adaptation in response to reductions in the reliability of sensory feedback^69–71^, speeding up of adaptation in response to uncertainty in the state of the individual or environment^28,67,71^, and even the optimal tuning of adaptation rates in individual subjects^30^.

Could this Bayesian framework also account for our results? The learning rate of a Kalman filter, and its steady-state properties depend on the ratio between two sources of variability: noise in the evolution of the generative process (e.g., perturbation) and the observation of trial-by-trial outcomes. If the brain were to interpret perturbation variability as an increase in observation noise, the Kalman framework would correctly predict that learning in the high-variance environment would proceed more slowly and saturate sooner than learning in the zero-variance environment. To fully capture our results, the Kalman framework would also require the brain to increase its estimate of process variability over time, in order to achieve increases in error sensitivity (i.e., Kalman gain) over the course of adaptation both in the zero-variance and high-variance environments (Fig. 4I).

However, there is one feature of our data that the standard Kalman filter cannot explain: variation in error sensitivity across different error magnitudes (Fig. 4A). While in both the zero-variance and high-variance groups error sensitivity declined as a function of error size^12,13,46,47^, perturbation variance affected only the sensitivity to small errors, not large errors. Eq. (1) explained this pattern; differences in perturbation variability led to changes in the consistency of small errors, but not large errors. It is unclear how to account for this phenomenon using a Kalman filter whose error sensitivity (i.e., Kalman gain) is independent of both error size as well as error history.

### Neural basis of implicit error sensitivity

Our finding that error consistency modulates the implicit component of adaptation raises important implications for the neural basis of error sensitivity. Implicit motor adaptation depends critically on the cerebellum^72–76^, where Purkinje cells learn to associate efference copies of motor commands with sensory consequences^77^. This learning is guided by sensory prediction errors, which are transmitted to the Purkinje cells via the inferior olive, resulting in complex spikes. Notably, plasticity in Purkinje cells exhibits both sensitivity to error, and forgetting. The response to error is determined by probability of complex spikes; in each Purkinje cell, the probability of complex spikes is greatest for a particular error vector^77,78^. Forgetting is present in the time-dependent retention of the plasticity caused by the complex spikes^42,79^, resulting in decay of plasticity with passage of time. Therefore, plasticity may saturate in the cerebellar cortex, limiting the total extent of adaptation.

Given these properties, how might perturbation variance alter the saturation of learning in the cerebellar cortex? One possibility is that the temporal consistency of complex spikes may alter the amount of plasticity experienced by each Purkinje cell. That is, when variance is low, errors of the same direction are likely to repeat, thus increasing the probability that the same population of Purkinje cells will experience multiple complex spikes in close temporal proximity. This theory makes the interesting prediction that the temporal proximity of complex spikes might modulate error sensitivity, thus altering the extent of adaptation. This idea remains to be tested.

## Methods

Here we describe the experiments and corresponding analysis reported in the main text. These include Experiments 1-7, as well as data reproduced by other sources including Fernandes and colleagues^19^, Robinson and colleagues^6^, and Kojima and colleagues^32^, and Kim and colleagues^13^.

### Participants

A total of 146 volunteers participated in our experiments. All experiments were approved by the Institutional Review Board at the Johns Hopkins School of Medicine.

### Apparatus

In Experiments 1-7, participants held the handle of a planar robotic arm (Fig. 1A) and made reaching movements to different target locations in the horizontal plane. The forearm was obscured from view by an opaque screen. An overhead projector displayed a small white cursor (diameter = 3mm) on the screen that tracked the motion of the hand. Throughout testing we recorded the position of the handle at submillimeter precision with a differential encoder. Data were recorded at 200 Hz.

### Visuomotor rotation

Experiments 1, 3, 4-7 followed a similar protocol. At the start of each trial, the participant brought their hand to a center starting position (circle with 1 cm diameter). After maintaining the hand within the start circle, a target circle (1 cm diameter) appeared in 1 of 4 positions (0°, 90°, 180°, and 270°) at a displacement of 8 cm from the starting circle. Participants then performed a “shooting” movement to move their hand briskly through the target. Each experiment consisted of epochs of 4 trials where each target was visited once in a pseudorandom order.

Participants were provided audiovisual feedback about their movement speed and accuracy. If a movement was too fast (duration < 75 ms) the target turned red. If a movement was too slow (duration > 325 ms) the target turned blue. If the movement was the correct speed, but the cursor missed the target, the target turned white. Successful movements (correct speed and placement) were rewarded with a point (total score displayed on-screen), an on-screen animation, and also a pleasing tone (1000 Hz). If the movement was unsuccessful, no point was awarded and a negative tone was played (200 Hz). Participants were instructed to obtain as many points as possible throughout the experimental session.

Once the hand reached the target, visual feedback of the cursor was removed, and a yellow marker was frozen on-screen to provide static feedback of the final hand position. At this point, participants were instructed to move their hand back to the starting position. The cursor continued to be hidden until the hand was moved within 2 cm of the starting circle. In most experiments, participants actively moved their hand back to the start position. However, in Experiments 3, 6, and 7 the robot assisted the subject if their hand had not returned to the start position after 1 second.

Movements were performed in one of three conditions: null trials, rotation trials, and no feedback trials. On null trials, veridical feedback of hand position was provided. On rotation trials, once the target appeared on screen, the on-screen cursor was rotated relative to the start position (Fig. 1A). Some rotation experiment terminated with a period of no feedback trials. On these trials, the subject cursor was hidden during the entire trial. No feedback was given regarding movement endpoint, accuracy, or timing.

As a measure of adaptation, we analyzed the reach angle on each trial. The reach angle was measured as the angle between the line segment connecting the start and target positions, and the line segment connecting the start and final hand position (defined as the point where the hand exceeded 95% of the target displacement). For analysis of reaching errors, we computed the same quantity, but for the final cursor position rather than the final hand position.

### Force field adaptation

In Experiment 2, participants were perturbed by a velocity-dependent force field (Fig. 1A), as opposed to a visuomotor rotation. At trial onset, a circular target (diameter= 1 cm) appeared in the workspace, coincident with a tone that cued subject movement. Participants then reached from the starting position to the target. The trial ended when the hand stopped within the target location. After stopping the hand within the target, feedback about movement duration was provided. If the preceding reach was too slow, the target turned blue and a low tone was played. If the reach was too fast, the target turned red and a low tone was played. If the reach fell within the desired movement interval (450-550 ms), the subject was rewarded with a point to their total score, an animation, and a pleasing tone (1000 Hz). Participants were instructed to obtain as many points as possible. After completing each outward reaching movement, participants were instructed to then bring their hand back to the starting position. This return movement was not rewarded and was always guided by a “channel” (see below).

As in the rotation experiments, the target appeared in 1 of 4 positions (0°, 90°, 180°, and 270°) at a displacement of 10 cm from the starting circle. Each experiment consisted of epochs of 4 trials where each target was visited once in a pseudorandom order. The experiment began with a set of null field trials (no perturbations from the robot). After this period, participants were exposed to a force field. The force field was a velocity-dependent curl field (Fig. 1A) in which the robot generated forces proportional and perpendicular to the velocity of the hand according to:

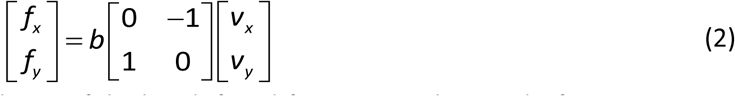

where *v*_*x*_ and *v*_*y*_ represent the x and y velocity of the hand, *f*_*x*_ and *f*_*y*_ represent the x and y force generated by the robot on the handle, and *b* represents the magnitude (and orientation) of the force field.

Subject reaching forces were measured on designated “channel” trials^36^ where the motion of the handle was restricted to a linear path connecting the start and target locations (Fig. 1A). To restrict hand motion to the straight-line channel trajectory, the robot applied perpendicular stiff spring-like forces with damping (stiffness = 6000 N/m, viscosity = 250 N-s/m). Reaching forces were measured on every 5^th^ epoch of movements with a cycle of 4 channel trials (one per target). In addition, the experiment terminated with a block of channel trials retention of the adapted state over trials.

Offline we isolated the perpendicular forces produced against the channel wall. We subtracted off the average force produced on channel trials during the baseline period. To measure adaptation, we calculated an adaptation index. The adaptation index represents the scaling factor relating the force produced on a given trial and the ideal force the subject would produce if they were fully adapted to the perturbation^27^. To calculate this scaling factor, we linearly regressed the ideal force timecourse (product of velocity and perturbation magnitude) onto the actual force timecourse.

In addition to analyzing the forces produced on channel trials, we also analyzed the trajectory of the hand on perturbation trials. From each trajectory we isolated a signed movement error, which we used to calculate the probability that an error switched sign from one trial to the next (Fig. 4C, Exp. 2). To calculate the movement error, we isolated the portion of each reaching movement between 20% and 90% of target displacement. Within this region we detected the maximum absolute error and treated this as the error magnitude. We signed this error according to whether the hand was to the left or right (or top or bottom) of the line connecting the start position and target position. To prevent minor overcompensations from being treated as movement errors, deviations that fell within 3 mm of the line connecting the start and target locations were not treated as errors. Using smaller thresholds of 1 or 2 mm did not qualitatively affect our results.

### Statistics

Statistical tests such as repeated measures ANOVA, two-way ANOVA, and mixed-ANOVA were carried out in IBM SPSS 25. In all cases we report the p-value, F-value, and 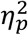 for each test. For post-hoc testing we employed t-tests with Bonferroni corrections. For these tests, we report the p-value and Cohen’s d as a measure of effect size. Our mixed-ANOVA contained a between-subjects factor and a within-subjects repeated measure. For the within-subjects repeated measure, data are binned within small windows defined by differences in error size. In the event that a participant is missing data within a bin (data are missing in approximately 13.2% of all bins), we replaced the missing data point with the mean of the appropriate distribution.

### Experiment 1

We tested how variance in the perturbation affected the total extent of visuomotor adaptation. The experiment started with 10 epochs (40 trials) of no perturbation. After this a perturbation period began that consisted of 60 rotation epochs (240 trials total). At the end of the perturbation period, retention of the visuomotor memory was tested in a series of 15 epochs (60 trials) of no feedback. To test the effect of perturbation variance on behavior, participants were divided into 1 of 2 groups. In the zero-variance group, participants (n=19) were exposed to a constant visuomotor rotation of 30°. In the high-variance group, participants (n=14) were exposed to a visuomotor rotation that changed on each trial. The rotation was sampled from a normal distribution with a mean of 30° and a standard deviation of 12°.

### Experiment 2

We found that perturbation variance reduced the total amount of adaptation in Experiment 1. To test if this impairment was a general property of sensorimotor adaptation, we tested another group of subjects with a force field. The experiment started with 10 epochs (40 trials) of no perturbation (2 of these epochs were channel trials). After this a perturbation period began that consisted of 75 epochs (300 trials, 20% were channel trials) of force field perturbations. At the end of the perturbation period, retention of the adapted state was tested in a series of 10 epochs (40 trials) of channel trial movements. To test the effect of perturbation variance on behavior, participants were divided into 1 of 2 groups. In the zero-variance group, participants (n=12) were exposed to a constant force field magnitude of 14 N-s/m. In the high-variance group, participants (n=13) were exposed to a force field magnitude that changed on each trial. The force field magnitude was sampled from a normal distribution with a mean of 14 N-s/m and a standard deviation of 6 N-s/m.

### Experiment 3

Inspection of the learning curves in Experiment 1 indicated that performance may not have completely saturated by the end of the perturbation period. Therefore, we repeated Experiment 1, but this time more than doubled the number of perturbation trials. The experiment started with 5 epochs (20 trials) of no perturbation. The following perturbation period consisted of 160 rotation epochs (640 trials). As in Experiment 1, participants were divided into a zero-variance group (n=10) and a high-variance group (n=10). Perturbation statistics remained identical to Experiment 1.

### Experiment 4

To determine if perturbation variance causally altered the total extent of adaptation, we designed a control experiment. In this experiment, participants started with a visuomotor rotation in the zero-variance condition, and then were exposed to the high-variance condition midway through the experiment. If variance causally determined the total amount of learning, we expected that asymptotic performance would decrease after the addition of variability to the perturbation. Participants (n=14) began the experiment with 5 epochs (20 trials) of null trials. After this, the zero-variance period started. Participants were exposed to either a CW or CCW visuomotor rotation of 30° for a total of 80 epochs (320 trials). At the end of this period, participants switched to a high-variance condition where the rotation was sampled on each trial from a normal distribution with a mean of 30° and a standard deviation of 12°. This period lasted for an additional 80 epochs (320 trials). Finally, the experiment concluded with 15 epochs (60 trials) of no feedback trials.

### Experiment 5

In Experiment 5, we suppressed implicit adaptation for the duration of the experiment, and measured the marginal effect of perturbation variability on the isolated explicit adaptation. To reduce implicit learning and isolate explicit strategy, we used experimental conditions that are well established to inhibit implicit learning. We removed all visual feedback of the cursor during the reach. Instead, only the terminal endpoint of the cursor was displayed, with a long delay of 1055 ms. In other words, visual feedback of the reach endpoint was shown approximately 1 second after the reach had ended. Delaying visual feedback has been shown to inhibit implicit recalibration of reach angle^25,26,39,40^. Apart from this change in feedback, all other details of the task were identical to Experiment 1.

### Experiment 6

In Experiment 6, we suppressed explicit adaptation for the duration of the experiment, and measured the marginal effect of perturbation variability on the isolated implicit adaptation. To isolate implicit adaptation, we limited the time participants had to prepare their movements. Limiting reaction time is known to suppress explicit strategy^22–24^. To limit reaction time, we instructed participants to begin their reaching movement as soon as possible, once the target was revealed. To enforce this, we limited the amount of time available for the participants to start their movement after the target location was shown. This upper bound on reaction time was set to either 225, 235, or 245 ms (taking into account screen delay). If the reaction time of the participant exceeded the desired upper bound, the participant was punished with a screen timeout after providing feedback of the movement endpoint. In addition, a low unpleasant tone (200 Hz) was played, and a message was provided on screen that read “React faster”. As in Experiment 1, participants were divided into a zero-variance perturbation group (n=13) and a high-variance group (n=12). All other details were identical to Experiment 1.

### Experiment 7

Sensorimotor adaptation is supported by both explicit strategy and implicit learning^10^. In Experiments 5 and 6, we isolated these systems so that each one alone adapted to the perturbation. In Experiment 7, we tested each simultaneously. The trial structure was equivalent to Experiment 1. Participants were placed into a zero-variance perturbation group (n=9; mean rotation of 30°, std. dev. of 0°) or a high-variance perturbation group (n=9; mean rotation of 30°, std. dev. of 12°). Participants performed 10 epochs of baseline no rotation trials, followed by 60 epochs (240 trials) of rotation trials.

After the last rotation epoch, the experiment was stopped briefly and the participants were provided with verbal instructions designed to isolate each participant’s implicit recalibration of reach angle^12,13,80^. Participants were told that for the next few trials there will be no cursor on the screen and no perturbation to the cursor position. Additionally, they were instructed to forget about the cursor, think only about their hand, and try to move their physical hand straight through the center of the target. After participants indicated that they understood the instructions, they performed one reaching movement to each of the 4 targets in a pseudorandom order without any visual feedback. The mean reach angle across the targets served as our measure of their final implicit reach angle (Fig. 2G, implicit, instruct). In addition, we subtracted this implicit reach angle from the mean reach angle measured over the last 10 epochs of the perturbation (prior to the verbal instruction) to estimate their explicit reach angle at the end of adaptation (Fig. 2G, explicit, instruct).

After this implicit probe period, we performed an additional test to directly assay each subject’s explicit re-aiming strategy. Each of the 4 targets was shown an additional time, with a ring of small white landmarks placed at an equal radial distance around the screen^9^. A total of 108 landmarks was used to uniformly cover the circle. Each landmark was labeled with a unique alphanumeric string. Participants were asked to report the nearest landmark that they were aiming towards at the end of the experiment in order to move the cursor through the target when the rotation was on. The mean angle reported across all 4 targets was calculated to provide an additional assay of explicit adaptation (Fig. 2G, explicit, clock). Explicit re-aiming is prone to erroneous selections where the participant mentally rotates the cursor in the wrong direction^23^ (errors of same magnitude, opposite sign). Therefore, for measurements where the participant reported an explicit angle in the opposite direction, we used its absolute value when calculating their explicit recalibration. Note that only 8 of the 9 participants in the high-variance group reported their aiming angles using this probe.

### State-space model of learning

After the experience of a movement error, humans and other animals change their behavior on future trials. In the absence of error, adapted behavior decays over time. Here we used a state-space model^81^ to capture this process of error-based learning. Here, the internal state of an individual *x*, changes from trials *n* to *n*+1 due to learning and forgetting.

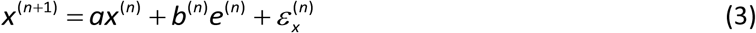

Forgetting is controlled by the retention factor *a*. The rate of learning is controlled by the error sensitivity *b*. Error sensitivity was modulated over time according to Eq. (1) in the main text. Learning and forgetting are stochastic processes affected by internal state noise *ε*_*x*_: a normal random variable with zero-mean and standard deviation of *σ*_*x*_.

While we cannot directly measure the internal state of an individual, we can measure their movements. The internal state *x* leads to a movement *y* according to:

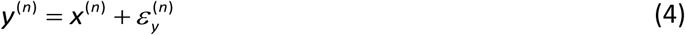

The desired movement is affected by execution noise, represented by *ε*_*y*_: a normal random variable with zero-mean and standard deviation of *σ*_*y*_. To complete the state-space model described by Eqs. 3 and 4, we must operationalize the value of an error, *e*. In sensorimotor adaptation, movement errors are determined both by motor output of the participant (*y*) and the size of the external perturbation (*r*):

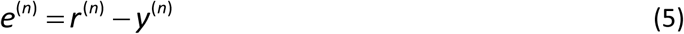

We used Eqs. (1,3-5) to produce motor output in Fig. 4. More details about the modulation of error sensitivity are provided below. In addition, we used Eqs. (3-5) with fixed error sensitivity to simulate the learning traces in Figs. 3A-C.

### Asymptotic properties of learning

State-space models of learning predict that performance saturates after prolonged exposure. This saturation is caused by a steady state condition where the amount of learning from error is exactly counterbalanced by the amount of forgetting (Fig. 3A). The steady state can be derived from Eqs. (3)-(5):

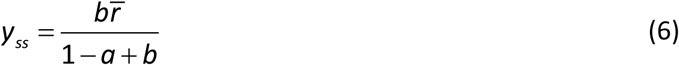

The formula for steady-state adaptation (*y*_ss_) shows that one’s learning extent depends on 3 factors: (1) error sensitivity *b*, (2) retention factor *a*, and (3) the mean of the perturbation 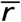. If there is no forgetting (*a* = 1), an individual will adapt completely to the mean of the perturbation. However, if retention is incomplete (*a* < 1), the steady state behavior (*y*_ss_) will always fall short of the mean of the perturbation, resulting in residual errors.

Eq. (6) is important for three reasons. (1) It demonstrates why the total extent of learning varies with a change in forgetting rate (Fig. 3B). (2) It demonstrates why the total extent of learning varies with a change in error sensitivity (Fig. 3C). (3) It demonstrates that the total amount of learning does not directly depend on variability in the perturbation, only the mean of the perturbation.

### Calculation of the retention factor

To determine if differences in learning extent were caused by a change in the rate of forgetting, we estimated the retention factor (*a*) of each participant. To do this, we quantified how behavior decayed during the error-free periods that terminated Experiments 1, 2, 4, and 6. During the error-free periods, trial errors were either hidden (no feedback condition in visuomotor rotation experiments) or fixed to zero (channel trials in the force field adaptation experiment). In the absence of error (*e*=0), our state-space model simplifies to exponential decay (omitting noise terms):

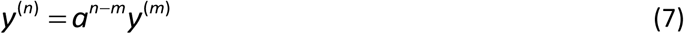

Eq. (7) relates motor output (*y*) on trial *n* of the error-free period to the initial motor behavior measured at the end of the adaptation period, *y*^(*m*)^. The term *n* – *m* represents the number of trials that elapsed from the start of the error-free period until the current trial *n*.

For visuomotor rotation experiments, we estimated the retention factor separately for each target by fitting Eq. (7) to subject behavior in the least-squares sense. We report the mean retention factor in Fig. 3F. For force field adaptation, we estimated a single retention factor, by first averaging the adaptation index across the 4 targets in each epoch, and then fitting Eq. (7) to the epoch-by-epoch behavior in the least-squares sense. In Fig. 3F, we converted this epoch-based retention factor to a trial-based retention factor by raising the epoch-based retention factor to the power of 1/4 (an epoch of 4 trials has 4 trial-by-trial decay events).

### Calculation of error sensitivity

Using Eq. (7), we found that changes in learning saturation were not caused by modulation of forgetting rates. Next, we determined how variability impacted error sensitivity (*b*), using its empirical definition:

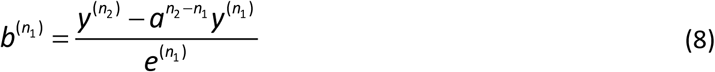

Eq. (8) determines the sensitivity to an error experienced on trial *n*_*1*_ when the participant visited a particular target T. This error sensitivity is equal to the change in behavior between two consecutive visits to target T, on trials *n*_*1*_ and *n*_2_ (i.e., there are no intervening trials where target T was visited) divided by the error that had been experienced on trial *n*_1_. In the numerator, we account for decay in the behavior by multiplying the behavior on trial *n*_1_ by a decay factor that accounted for the number of intervening trials between trials *n*_1_ and *n*_2_. For each target, we used the specific retention factor estimated for that target with Eq. (7).

We used Eq. (8) to calculate error sensitivity for all of our visuomotor rotation experiments. When reporting error sensitivity, we averaged across the four targets (Figs. 3G, 4A, 4I, 4J). In some cases (Fig. 3G) we collapsed trial-by-trial measurements of error sensitivity across all trials and all errors. In other cases, we calculated the change in error sensitivity over different periods of training. For Fig. 4J, we measured the change in sensitivity from the beginning (epochs 1-10) to the end (epochs 49-59) of the perturbation block in Exp. 6 (implicit only). To remove outliers, we identified error sensitivity estimates that deviated from the population median by over 3 median absolute deviations. We did this within windows of 10 epochs. This procedure was also used to compute the timecourse in Fig. 4I.

In Fig. 4A, we calculated error sensitivity for errors of different sizes combining together data from Exps. 1, 4, and 6. We divided up the error space into bins of small errors (5-14°), medium errors (14-22°), and large errors (22-30°). To prevent noisy estimates of error sensitivity from populating each bin, we added a subject to a bin contingent on them at least having 12 trials (5% of the total number of adaptation trials) for which an error was experienced in the corresponding range). We did not consider errors smaller than 5° because the empirical estimator in Eq. (8) becomes unstable for small error sizes.

For force field adaptation, we could not empirically estimate error sensitivity, as this approach requires the measurement of forces directly before and after the experience of an error. However, in reality, forces are measured only on infrequent channel trials, making such an empirical calculation impossible. For this reason, we used a model-based approach to measure error sensitivity (Fig. 3G, Exp. 2). We fit our state-space model Eqs. (3-5) to single subject data in the least-squares sense, over the last 5 channel trial epochs of the adaptation period. To do this, we needed to describe four states of learning (one for each target). We describe multitarget state-space models in more detail in an earlier work^81^. As a brief summary, we modeled our multitarget experiment by applying Eqs. (3-5) separately for each target. On any given trial, the state corresponding to the relevant target learned from the error on that trial. The other three states exhibited only decay on that trial. We described the perturbation *r* in terms of the force field magnitude on that trial (14 N-s/m was considered a perturbation of unit 1 in the model). Using this framework, we found the error sensitivity that minimized the squared difference between our model simulation and participant behavior.

### Decaying memory of errors model

To account for the relationship between error sensitivity and error consistency (Fig. 4A) we adapted the memory of errors model proposed by Herzfeld and colleagues^31^. This model uses a simple normative framework. When the errors on trial *n* and trial *n*+1 have the same sign (a consistent error), this signals that the brain under-corrected for the first error (Fig. 4B, left). Therefore, the brain should increase its sensitivity to the initial error. On the other hand, when the errors on trials *n* and *n*+1 have opposite signs (an inconsistent error), this signals that the brain over-corrected for the first error (Fig. 4B, right). Therefore, the brain should decrease its sensitivity to the initial error. These rules are encapsulated by the right-most term of Eq. (1).

The right-most term in Eq. (1) alone accounts for a rich set of behavioral phenomena including savings and meta-learning^31^. However, its ability to describe saturation of learning is limited by its lack of decay. Without trial-based decay in error sensitivity, common experimental conditions prevent the model from reaching a saturation point. For this reason, our adapted memory of errors model (Eq. (1)) includes a term for learning, controlled by the parameter β, and a term for decay, controlled by the parameter α.

The combination of α, β, and trial-to-trial error consistency determine how error sensitivity changes over time. Critically, error sensitivity changes locally, that is, only errors near that experienced on trial *n* will experience an upregulation or downregulation in sensitivity. To enforce this, the ***c*** vector in Eq. (1) has all but one entry equal to zero. The vector contains a single value of one, at the index corresponding to an error window containing the error experienced on trial *n*. For our model predictions in Fig. 4, we spaced non-overlapping error windows between errors of -100° and 100°, each with a width of 5°. The term **Δ*b*** is a vector whose elements represents the change in error sensitivity within each 5° bin. On any given trial, the error sensitivity of the learner was obtained through:

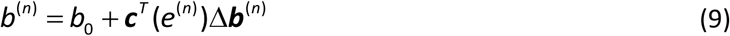

Here *b*_*0*_ represents the baseline error sensitivity of the system. Altogether, Eqs. (2-5) describe our state-space model whose error sensitivity is updated trial-by-trial according to Eqs. (1) and (9).

In Fig. 4, we fit our decaying memory of errors model to the implicit-only behavior measured under reaction time restrictions in Experiment 6. We fit the two free parameters, α and β, to the mean reach angles in the least-squares sense. Both the zero-variance and the high-variance groups were fit at the same time with the same parameter set. For the fitting process, we fixed all other model parameters to empirical measurements. For the initial error sensitivity *b*_*0*_, we used the median initial error sensitivity (0.037) measured across the zero-variance and high-variance groups in Experiment 6. For the retention factor, we again combined both groups, converted trial-based retention factors to epoch-based retention factors, averaged the retention factor across all 4 targets, removed any retention factors greater than one, and then calculated the midpoint of the resulting distribution. This yielded an epoch-by-epoch retention factor of 0.9134.

To identify the optimal values of α and β, we used the following grid-search procedure across all pairwise combinations of α (300 values spaced evenly between 0.95 and 0.995) and β (300 values spaced evenly between 0.01 and 0.1). For any given pair of α and β, we used Eq. (1) to predict changes in error sensitivity within each 5° bin. We did this for the exact error sequence measured in each individual participant. That is, for each participant, we used their error sequence, α, and β to predict how error sensitivity should vary as function of trial and error size. This process yielded a separate error sensitivity for each error size. These multiple timeseries were collapsed into one, by selecting the error sensitivity corresponding to the error experienced on the appropriate trial. For example, if on trial *m* the participant experienced an error in bin *b*, the collapsed timecourse used the predicted error sensitivity in bin *b* on trial *m*. We did this for all 4 targets separately, and then averaged the predicted timecourses across the targets. Finally, we then averaged across participants. In this way, we used the actual error consistency in each participant along with Eq. (1) to predict error sensitivity as a function of trial. The noisy traces in Fig. 4F show the mean predicted error sensitivity for the optimal parameter set.

Next, we used this mean predicted error sensitivity to simulate the state-space model specified by Eqs. (3-5). In other words, we simulated Eqs. (3-5) varying error sensitivity from one trial to the next according to the predicted trace obtained from Eq. (1). Note that this process would tend to yield a noisy adaptation profile, as the underlying estimates of error sensitivity were noisy (see noisy traces in Fig. 4F). Therefore, we used a smoothed version of these estimates for our simulation. These smoothed estimates are depicted by the blue lines in Fig. 4F. To produce these smoothed traces, we used a piecewise fit to the data. We divided the error sensitivity trace into two parts (for the zero-variance perturbation, these parts were divided on epoch 20; for the high-variance perturbation, these parts were divided on epoch 7). For the first part, we fit an exponential function that minimized the squared error between the empirical fit and measured error sensitivity. This fit was constrained to begin at *b*_*0*_, and terminate continuously with the smoothed fit to the second part of the data. For the second part of the data we fit a cubic smoothing spline using the *csaps* function in MATLAB R2019a with a roughness measure of 0.0003.

Altogether, for any set of α and β, this yielded a mean predicted behavior. We identified the α and β that minimized the squared error between the model predictions and the measured behavior across the zero-variance and high-variance groups. This yielded α = 0.9568 and β = 0.0558. Using these parameters, we not only simulated the expected error sensitivity timecourse in Fig. 4F, but also the corresponding learning curve in Fig. 4G (model). In Fig. 4J we report the change in error sensitivity predicted across the zero-variance and high-variance group. For this, we calculated the change in predicted implicit error sensitivity from the first 10 epochs to the last 10 epochs in Fig. 4F.

### Fernandes and colleagues (2012)

In Fig. 1B, we reference earlier work from a study by Fernandes and colleagues^19^. Briefly, participants (n=16) made a center-out reaching movement to a target. After the reach ended, participants were shown the final location of the right index finger. Participants performed three experimental blocks. Each block had the same general structure. At the start of the block, participants made 40 reaching movements to 8 different targets (5 for each target) with continuous visual feedback of the cursor. Next, participants made an additional 80 reaching movements to 8 different targets (10 for each target) using only endpoint feedback of the cursor position. After this baseline period, a single target position was selected, and 240 reaching movements were performed under the influence of a visuomotor rotation. The visuomotor rotation was sampled on each trial from a normal distribution with a mean of 30° and a standard deviation of either 0°, 4°, or 12°. The block ended in a set of 160 generalization trials that are not relevant to the current study. The experiment had a within-subject design. Each participant was exposed to all three perturbation variances, but in a random order. The orientation of the rotation (CW or CCW) was randomly chosen on each block. In addition, the target selected during the adaptation period was randomly chosen from 1 of the 4 diagonal targets on each block.

### Kim and colleagues (2018)

We compared the implicit learning measured under reaction time restrictions in Experiment 6, to the implicit learning measured under the constant error-clamp conditions reported by Kim and colleagues^13^. Specifically, we calculated the standard deviation of the terminal amount of implicit learning reported under both conditions. For Kim and colleagues, we visually inspected Fig. 2b of the corresponding manuscript in Adobe Illustrator to obtain the asymptotic implicit hand angle for the 1.75° clamp group, the 3.5° clamp group, and the 15° clamp group. We collapsed participants across groups and then calculated the standard deviation of the resulting distribution. For our data, we considered the zero-variance group in Experiment 6. We calculated the mean reaching angle on the last 2 cycles of the rotation period. We used 2 cycles as this would equal 8 total trials, which matches the number of trials included in the constant error-clamp measure. We then calculated the standard deviation of terminal implicit angles across all participants. For our data, the standard deviation was 3.17°. For Kim and colleagues^13^ the standard deviation was 9.59°, representing an increase of approximately 300% over our measure of implicit learning.

### Robinson and colleagues (2003)

Robinson and colleagues^6^ adapted monkeys to a saccadic perturbation, where the error on every trial was fixed to -1° independent of the monkey’s motor output (Fig. 5A). Critically, despite the fact that error never decreased, learning still reached a saturation point. To reach this steady-state, sensitivity to error must also reach an asymptotic limit. How does the memory of errors account for this limit?

Here we fit two variants of the memory of errors model to these constant error-clamp data, shown in the middle inset of Fig. 5B. One of these models assumed that error sensitivity did not decay from one trial to the next (α=1). The other model allowed error sensitivity to decay (α<1). To fit these models to the measured data, we extracted behavior from the original manuscript using the GRABIT routine in MATLAB R2018a. For our simulations, we set the initial error sensitivity to 0.005 and used a retention factor of 0.98. We divided up the error sensitivity bins in Eq. (1) into 100 windows spaced evenly between errors of -6 and 6°. Also, we simulated deterministic behavior by setting σ_x_ from Eq. (3) and s_y_ from Eq. (4) both equal to 0°.

To fit the decaying model and decay-free model to the behavior, we used *fmincon* in MATLAB R2018a to identify the parameter set that minimized the sum of squared error between the model predictions and measured behavior. We predicted behavior using the state-space model specified by Eqs. (3-5) with an error sensitivity that varied according to Eq. (1). To account for the initial bias in saccade gain, we subtracted off a gain of 0.133 from the behavior predicted by our state-space model. For each model, we performed 100 iterations of *fmincon* each time varying the parameter set used to seed the algorithm. For the decay-free model (α=1), the optimal value of β was 8.163×10^−5^. For the decaying model, the optimal parameter set was α=0.9883 and β=0.0006. The behavior predicted by each model is shown in the middle inset of Fig. 5. The corresponding error sensitivity is shown at the bottom of Fig. 5.

### Kojima and colleagues (2004)

Kojima and colleagues^32^ exposed monkeys to a 3.5° visual perturbation, then a -3.5° perturbation, followed by re-exposure to the original 3.5° perturbation (Fig. 5B, no zero-error period). This paradigm elicited savings, a faster rate of re-learning that is linked to increases in error sensitivity^29,31,48,49^ (Fig. 5B, compare initial rates of learning denoted by the linear regression lines in the middle inset). However, when a long period of no perturbation trials was inserted after washout (Fig. 5C, zero-error period), no savings was observed (Fig. 5C, compare initial rates of learning denoted by the linear regression lines). These data provide clear evidence that error sensitivity decays over long time scales. How do the decay (α<1) and decay-free (α=1) variants of the memory of errors model account for the dissolution of savings with extended washout?

Data from their original manuscript is reproduced in Figs. 5B-D. Here we contrast the predictions of the decay-free and decaying model. We simulated these models in the short washout paradigm in Fig. 5B. For the short washout paradigm, we simulated 750 trials of a 3.5° gain-up perturbation, followed by 417 trials of a -3.5° gain-down perturbation, followed by 750 trials of the 3.5° gain-up perturbation. We chose 417 trials for the gain down perturbation because at this trial behavior reached baseline saccade amplitude. For the long washout paradigm in Fig. 5C, we simulated the same schedule, only adding 780 trials of zero perturbation trials prior to re-exposure to the 3.5° perturbation. We chose 780 trials to match the paradigm reported by Kojima and colleagues^32^.

To simulate each model, we used a retention factor of 1, an initial error sensitivity of 8.6×10^−4^, and 30 error sensitivity bins (Eq. (1)) spaced evenly between errors of -6 and 6°. For both the decay and decay-free models, we used β=1.25×10^−5^. We selected these parameters so that the model predictions matched the early learning rates reported in the original manuscript. That is, the slope over the first 150 trials of the first and second exposures to the gain-up perturbation was equal to 4×10^−4^ and 6.9×10^−4^ °/trial, respectively. For the no-decay model, α was set to 1 for the entirely of the simulation. For simplicity of comparison, we matched the behavior of the decaying model to the decay-free by starting with these same parameters. However, during the zero-error period, we set the α parameter to 0.989 for the decaying model. This value was selected from our main result in Fig. 4 (here the epoch-by-epoch decay parameter was equal to 0.9568, and so we raised it to the 0.25 power to obtain a trial-by-trial decay parameter). Finally, we simulated stochastic output of the decay and no-decay models, setting s_x_ from Eq. (3) equal to 0°, and s_y_ from Eq. (4) equal to 0.2°.

The behaviors predicted by the decay and no-decay models are shown in Figs. 5B and 5C, at bottom. These curves represent the mean behavior predicted across 100,000 simulations of the state-space model. We quantified savings similar to the original manuscript by Kojima and colleagues^32^, using linear regression. Here, we linearly regressed the simulated behavior onto the trial counts over the periods designated by “i", “ii”, and “iii” in Figs. 5B and C. These periods represent the first 150 trials of the perturbation. Then we calculated the percent change in rate from “i” to “ii” (for the short washout experiment) and “i” to “iii” for the long washout experiment (Fig. 5D). We compared these predicted values for the decaying model and decay-free model, to the empirical measurements reported in the original manuscript (these values are shown next to the regression lines in Figs. 5B and 5C, and are represented by the black bars in Fig. 5D).

## Acknowledgements

This work was supported by grants from the National Institutes of Health (R01NS078311, F32NS095706), the National Science Foundation (CNS-1714623), the Cambridge Trust, the Rutherford Foundation, and a travel grant from the Boehringer Ingelheim Fonds. Additionally, we would like to thank Hugo Fernandes and Konrad Kording for so graciously compiling and sharing their data with us. Finally, we would like to recognize the Summer School in Computational Sensory-Motor Neuroscience (CoSMo) and its organizers (Gunnar Blohm, Konrad Kording, and Paul Schrater), for giving us the opportunity to learn and develop the original idea for this work.

## References

1. Donchin, O. et al. Cerebellar regions involved in adaptation to force field and visuomotor perturbation. J. Neurophysiol. 107, 134–147 (2012).

2. Brashers-Krug, T., Shadmehr, R. & Bizzi, E. Consolidation in human motor memory. Nature 382, 252–255 (1996).

3. Tseng, Y.-W., Diedrichsen, J., Krakauer, J. W., Shadmehr, R. & Bastian, A. J. Sensory prediction errors drive cerebellum-dependent adaptation of reaching. J. Neurophysiol. 98, 54–62 (2007).

4. Krakauer, J. W., Pine, Z. M., Ghilardi, M. F. & Ghez, C. Learning of visuomotor transformations for vectorial planning of reaching trajectories. J. Neurosci. 20, 8916–8924 (2000).

5. Ethier, V., Zee, D. S. & Shadmehr, R. Spontaneous Recovery of Motor Memory During Saccade Adaptation. J. Neurophysiol. 99, 2577–2583 (2008).

6. Robinson, F. R., Noto, C. T. & Bevans, S. E. Effect of Visual Error Size on Saccade Adaptation in Monkey. J. Neurophysiol. 90, 1235–1244 (2003).

7. Malone, L. A., Vasudevan, E. V. L. & Bastian, A. J. Motor Adaptation Training for Faster Relearning. J. Neurosci. 31, 15136–15143 (2011).

8. Shadmehr, R., Brandt, J. & Corkin, S. Time-dependent motor memory processes in amnesic subjects. J. Neurophysiol. 80, 1590–1597 (1998).

9. McDougle, S. D., Bond, K. M. & Taylor, J. A. Explicit and Implicit Processes Constitute the Fast and Slow Processes of Sensorimotor Learning. J. Neurosci. 35, 9568–9579 (2015).

10. Taylor, J. A., Krakauer, J. W. & Ivry, R. B. Explicit and Implicit Contributions to Learning in a Sensorimotor Adaptation Task. J. Neurosci. 34, 3023–3032 (2014).

11. Taylor, J. A. & Ivry, R. B. Flexible cognitive strategies during motor learning. PLoS Comput. Biol. 7, e1001096 (2011).

12. Morehead, J. R., Taylor, J. A., Parvin, D. E. & Ivry, R. B. Characteristics of Implicit Sensorimotor Adaptation Revealed by Task-irrelevant Clamped Feedback. J. Cogn. Neurosci. 29, 1061–1074 (2017).

13. Kim, H. E., Morehead, J. R., Parvin, D. E., Moazzezi, R. & Ivry, R. B. Invariant errors reveal limitations in motor correction rather than constraints on error sensitivity. Commun. Biol. 1, 19 (2018).

14. Hegele, M. & Heuer, H. The impact of augmented information on visuo-motor adaptation in younger and older adults. PLoS One 5, e12071–e12071 (2010).

15. Heuer, H. & Hegele, M. Adaptation to visuomotor rotations in younger and older adults. Psychol. Aging 23, 190–202 (2008).

16. Vandevoorde, K. & Orban de Xivry, J.-J. Internal model recalibration does not deteriorate with age while motor adaptation does. Neurobiol. Aging 80, 138–153 (2019).

17. Vaswani, P. A. et al. Persistent Residual Errors in Motor Adaptation Tasks: Reversion to Baseline and Exploratory Escape. J. Neurosci. 35, 6969–6977 (2015).

18. Langsdorf, L., Maresch, J., Hegele, M., McDougle, S. D. & Schween, R. Prolonged reaction times eliminate residual errors in visuomotor adaptation. bioRxiv (2019). doi:10.1101/2019.12.26.888941

19. Fernandes, H. L., Stevenson, I. H. & Kording, K. P. Generalization of stochastic visuomotor rotations. PLoS One 7, e43016 (2012).

20. Therrien, A. S., Wolpert, D. M. & Bastian, A. J. Increasing Motor Noise Impairs Reinforcement Learning in Healthy Individuals. eNeuro 5, (2018).

21. Havermann, K. & Lappe, M. The Influence of the Consistency of Postsaccadic Visual Errors on Saccadic Adaptation. J. Neurophysiol. 103, 3302–3310 (2010).

22. Fernandez-Ruiz, J., Wong, W., Armstrong, I. T. & Flanagan, J. R. Relation between reaction time and reach errors during visuomotor adaptation. Behav. Brain Res. 219, 8–14 (2011).

23. McDougle, S. D. & Taylor, J. A. Dissociable cognitive strategies for sensorimotor learning. Nat. Commun. 10, 40 (2019).

24. Leow, L.-A., Marinovic, W., de Rugy, A. & Carroll, T. J. Task errors drive memories that improve sensorimotor adaptation. J. Neurosci. (2020). doi:10.1523/JNEUROSCI.1506-19.2020

25. Held, R., Efstathiou, A. & Greene, M. Adaptation to displaced and delayed visual feedback from the hand. J. Exp. Psychol. 72, 887–891 (1966).

26. Schween, R. & Hegele, M. Feedback delay attenuates implicit but facilitates explicit adjustments to a visuomotor rotation. Neurobiol. Learn. Mem. 140, 124–133 (2017).

27. Smith, M. A., Ghazizadeh, A. & Shadmehr, R. Interacting adaptive processes with different timescales underlie short-term motor learning. PLoS Biol. 4, e179 (2006).

28. Kording, K. P., Tenenbaum, J. B. & Shadmehr, R. The dynamics of memory as a consequence of optimal adaptation to a changing body. Nat. Neurosci. 10, 779–786 (2007).

29. Coltman, S. K., Cashaback, J. G. A. & Gribble, P. L. Both fast and slow learning processes contribute to savings following sensorimotor adaptation. J. Neurophysiol. 121, 1575–1583 (2019).

30. van der Vliet, R. et al. Individual Differences in Motor Noise and Adaptation Rate Are Optimally Related. eNeuro 5, (2018).

31. Herzfeld, D. J., Vaswani, P. A., Marko, M. K. & Shadmehr, R. A memory of errors in sensorimotor learning. Science (80-.). 345, 1349–1353 (2014).

32. Kojima, Y., Iwamoto, Y. & Yoshida, K. Memory of Learning Facilitates Saccadic Adaptation in the Monkey. J. Neurosci. 24, 7531–7539 (2004).

33. Kitago, T., Ryan, S. L., Mazzoni, P., Krakauer, J. W. & Haith, A. M. Unlearning versus savings in visuomotor adaptation: comparing effects of washout, passage of time, and removal of errors on motor memory. Front. Hum. Neurosci. 7, 307 (2013).

34. Leow, L.-A., de Rugy, A., Marinovic, W., Riek, S. & Carroll, T. J. Savings for visuomotor adaptation require prior history of error, not prior repetition of successful actions. J. Neurophysiol. 116, 1603–1614 (2016).

35. Sing, G. C. & Smith, M. A. Reduction in learning rates associated with anterograde interference results from interactions between different timescales in motor adaptation. PLoS Comput. Biol. 6, e1000893 (2010).

36. Scheidt, R. A., Reinkensmeyer, D. J., Conditt, M., Rymer, W. Z. & Mussa-ivaldi, F. A. Persistence of motor adaptation during constrained, multi-joint, arm movements. J. Neurophysiol. 84, 853–862 (2000).

37. Mazzoni, P. & Krakauer, J. W. An implicit plan overrides an explicit strategy during visuomotor adaptation. J. Neurosci. 26, 3642–3645 (2006).

38. Hwang, E. J., Smith, M. A. & Shadmehr, R. Dissociable effects of the implicit and explicit memory systems on learning control of reaching. Exp. brain Res. 173, 425–437 (2006).

39. Schween, R., Taube, W., Gollhofer, A. & Leukel, C. Online and post-trial feedback differentially affect implicit adaptation to a visuomotor rotation. Exp. brain Res. 232, 3007–3013 (2014).

40. Brudner, S. N., Kethidi, N., Graeupner, D., Ivry, R. B. & Taylor, J. A. Delayed feedback during sensorimotor learning selectively disrupts adaptation but not strategy use. J. Neurophysiol. 115, 1499–1511 (2016).

41. Ekerot, C. F. & Kano, M. Stimulation parameters influencing climbing fibre induced long-term depression of parallel fibre synapses. Neurosci. Res. 6, 264–268 (1989).

42. Herzfeld, D. J., Kojima, Y., Soetedjo, R. & Shadmehr, R. Encoding of error and learning to correct that error by the Purkinje cells of the cerebellum. Nat. Neurosci. 21, 736–743 (2018).

43. Haith, A. M., Huberdeau, D. M. & Krakauer, J. W. The Influence of Movement Preparation Time on the Expression of Visuomotor Learning and Savings. J. Neurosci. 35, 5109–5117 (2015).

44. Leow, L.-A., Gunn, R., Marinovic, W. & Carroll, T. J. Estimating the implicit component of visuomotor rotation learning by constraining movement preparation time. J. Neurophysiol. 118, 666–676 (2017).

45. Maresch, J., Werner, S. & Donchin, O. Methods matter: your measures of explicit and implicit processes in visuomotor adaptation affect your results. bioRxiv (2020). doi:10.1101/702290

46. Marko, M. K., Haith, A. M., Harran, M. D. & Shadmehr, R. Sensitivity to prediction error in reach adaptation. J Neurophysiol 108, 1752–1763 (2012).

47. Wei, K. & Kording, K. Relevance of error: what drives motor adaptation? J. Neurophysiol. 101, 655–664 (2009).

48. Zarahn, E., Weston, G. D., Liang, J., Mazzoni, P. & Krakauer, J. W. Explaining Savings for Visuomotor Adaptation: Linear Time-Invariant State-Space Models Are Not Sufficient. J. Neurophysiol. 100, 2537–2548 (2008).

49. Mawase, F., Shmuelof, L., Bar-Haim, S. & Karniel, A. Savings in locomotor adaptation explained by changes in learning parameters following initial adaptation. J. Neurophysiol. 111, 1444–1454 (2014).

50. Yin, C. & Wei, K. Savings in sensorimotor adaptation without an explicit strategy. J. Neurophysiol. 123, 1180–1192 (2020).

51. Wilterson, S. A. & Taylor, J. A. Implicit visuomotor adaptation remains limited after several days of training. bioRxiv (2019). doi:10.1101/711598

52. Morehead, J. R., Qasim, S. E., Crossley, M. J. & Ivry, R. Savings upon Re-Aiming in Visuomotor Adaptation. J. Neurosci. 35, 14386–14396 (2015).

53. Bond, K. M. & Taylor, J. A. Flexible explicit but rigid implicit learning in a visuomotor adaptation task. J. Neurophysiol. 113, 3836–3849 (2015).

54. Avraham, G., Keizman, M. & Shmuelof, L. Environmental Consistency Modulation of Error Sensitivity During Motor Adaptation is Explicitly Controlled. J. Neurophysiol. (2019). doi:10.1152/jn.00080.2019

55. Huberdeau, D. M., Haith, A. M. & Krakauer, J. W. Formation of a long-term memory for visuomotor adaptation following only a few trials of practice. J. Neurophysiol. 114, 969–977 (2015).

56. Neville, K.-M. & Cressman, E. K. The influence of awareness on explicit and implicit contributions to visuomotor adaptation over time. Exp. Brain Res. 236, 2047–2059 (2018).

57. Miyamoto, Y. R., Wang, S. & Smith, M. A. Implicit adaptation compensates for erratic explicit strategy in human motor learning. Nat. Neurosci. 23, 443–455 (2020).

58. Gonzalez Castro, L. N., Hadjiosif, A. M., Hemphill, M. A. & Smith, M. A. Environmental Consistency Determines the Rate of Motor Adaptation. Curr. Biol. 24, 1050–1061 (2014).

59. Smith, M. A. & Shadmehr, R. Modulation of the rate of error-dependent learning by statistical properties of the task. in Advances in Computational Motor Control (2004).

60. Lerner, G. et al. The origins of anterograde interference in visuomotor adaptation. bioRxiv 593996 (2019). doi:10.1101/593996

61. Thoroughman, K. & Shadmehr, R. Learning of action through adaptive combination of motor primitives. Nature 407, 742–7 (2000).

62. van der Kooij, K., Brenner, E., van Beers, R. J. & Smeets, J. B. J. Visuomotor adaptation: how forgetting keeps us conservative. PLoS One 10, e0117901 (2015).

63. Robinson, F. R., Soetedjo, R. & Noto, C. Distinct short-term and long-term adaptation to reduce saccade size in monkey. J. Neurophysiol. 96, 1030–1041 (2006).

64. Kojima, Y. & Soetedjo, R. Change in sensitivity to visual error in superior colliculus during saccade adaptation. Sci. Rep. 7, 9566 (2017).

65. Kojima, Y. & Soetedjo, R. Elimination of the error signal in the superior colliculus impairs saccade motor learning. Proc. Natl. Acad. Sci. 115, E8987--E8995 (2018).

66. Huang, V. S., Haith, A., Mazzoni, P. & Krakauer, J. W. Rethinking motor learning and savings in adaptation paradigms: model-free memory for successful actions combines with internal models. Neuron 70, 787–801 (2011).

67. Baddeley, R. J., Ingram, H. A. & Miall, R. C. System identification applied to a visuomotor task: near-optimal human performance in a noisy changing task. J. Neurosci. 23, 3066–3075 (2003).

68. Kalman, R. A New Approach to Linear Filtering and Prediction Problems. ASME J. Basic Eng. 34–45 (1960).

69. Burge, J., Ernst, M. O. & Banks, M. S. The statistical determinants of adaptation rate in human reaching. J. Vis. 8, 1–19 (2008).

70. van Beers, R. J. How does our motor system determine its learning rate? PLoS One 7, e49373–e49373 (2012).

71. Wei, K. & Körding, K. Uncertainty of feedback and state estimation determines the speed of motor adaptation. Front. Comput. Neurosci. 4, 11 (2010).

72. Xu-Wilson, M., Chen-Harris, H., Zee, D. S. & Shadmehr, R. Cerebellar Contributions to Adaptive Control of Saccades in Humans. J. Neurosci. 29, 12930–12939 (2009).

73. Galea, J. M., Vazquez, A., Pasricha, N., Orban De Xivry, J. J. & Celnik, P. Dissociating the roles of the cerebellum and motor cortex during adaptive learning: The motor cortex retains what the cerebellum learns. Cereb. Cortex 21, 1761–1770 (2011).

74. Herzfeld, D. J. et al. Contributions of the cerebellum and the motor cortex to acquisition and retention of motor memories. Neuroimage 98, 147–158 (2014).

75. Hanajima, R. et al. Modulation of error-sensitivity during a prism adaptation task in people with cerebellar degeneration. J. Neurophysiol. 114, 2460–2471 (2015).

76. Kim, S., Ogawa, K., Lv, J., Schweighofer, N. & Imamizu, H. Neural Substrates Related to Motor Memory with Multiple Timescales in Sensorimotor Adaptation. PLoS Biol. 13, e1002312 (2015).

77. Herzfeld, D. J., Kojima, Y., Soetedjo, R. & Shadmehr, R. Encoding of action by the Purkinje cells of the cerebellum. Nature 526, 439–442 (2015).

78. Soetedjo, R., Kojima, Y. & Fuchs, A. F. Complex spike activity in the oculomotor vermis of the cerebellum: a vectorial error signal for saccade motor learning? J. Neurophysiol. 100, 1949–1966 (2008).

79. Yang, Y. & Lisberger, S. G. Role of plasticity at different sites across the time course of cerebellar motor learning. J. Neurosci. 34, 7077–90 (2014).

80. Kim, H. E., Parvin, D. E. & Ivry, R. B. The influence of task outcome on implicit motor learning. Elife 8, e39882 (2019).

81. Albert, S. T. & Shadmehr, R. Estimating properties of the fast and slow adaptive processes during sensorimotor adaptation. J. Neurophysiol. 119, 1367–1393 (2018).

